# Pattern-Driven Navigation in 2D Multiscale Visualizations with Scalable Insets

**DOI:** 10.1101/301036

**Authors:** Fritz Lekschas, Michael Behrisch, Benjamin Bach, Peter Kerpedjiev, Nils Gehlenborg, Hanspeter Pfister

## Abstract

We present *Scalable Insets*, a technique for interactively exploring and navigating large numbers of annotated patterns in multiscale visualizations such as gigapixel images, matrices, or maps. Exploration of many but sparsely-distributed patterns in multiscale visualizations is challenging as visual representations change across zoom levels, context and navigational cues get lost upon zooming, and navigation is time consuming. Our technique visualizes annotated patterns too small to be identifiable at certain zoom levels using insets, i.e., magnified thumbnail views of the annotated patterns. Insets support users in searching, comparing, and contextualizing patterns while reducing the amount of navigation needed. They are dynamically placed either within the viewport or along the boundary of the viewport to offer a compromise between locality and context preservation. Annotated patterns are interactively clustered by location and type. They are visually represented as an aggregated inset to provide scalable exploration within a single viewport. In a controlled user study with 18 participants, we found that Scalable Insets can speed up visual search and improve the accuracy of pattern comparison at the cost of slower frequency estimation compared to a baseline technique. A second study with 6 experts in the field of genomics showed that Scalable Insets is easy to learn and provides first insights into how Scalable Insets can be applied in an open-ended data exploration scenario.

## 1 Introduction

Many large datasets, such as gigapixel images, geographic maps, or networks, require exploration of annotations at different levels of detail. We call an annotation a region in the visualization that contains some visual patterns (called *annotated pattern* henceforth) as seen in Figure 1. These annotated patterns can either be generated by users or derived computationally. However, annotated patterns are often magnitudes smaller than the overview and too small to be identifiable, also known as “desert fog” [33]. This makes exploring, searching, comparing, and contextualizing challenging, as considerable navigation is needed to overcome the lack of overview or detail.

**Figure 1:**
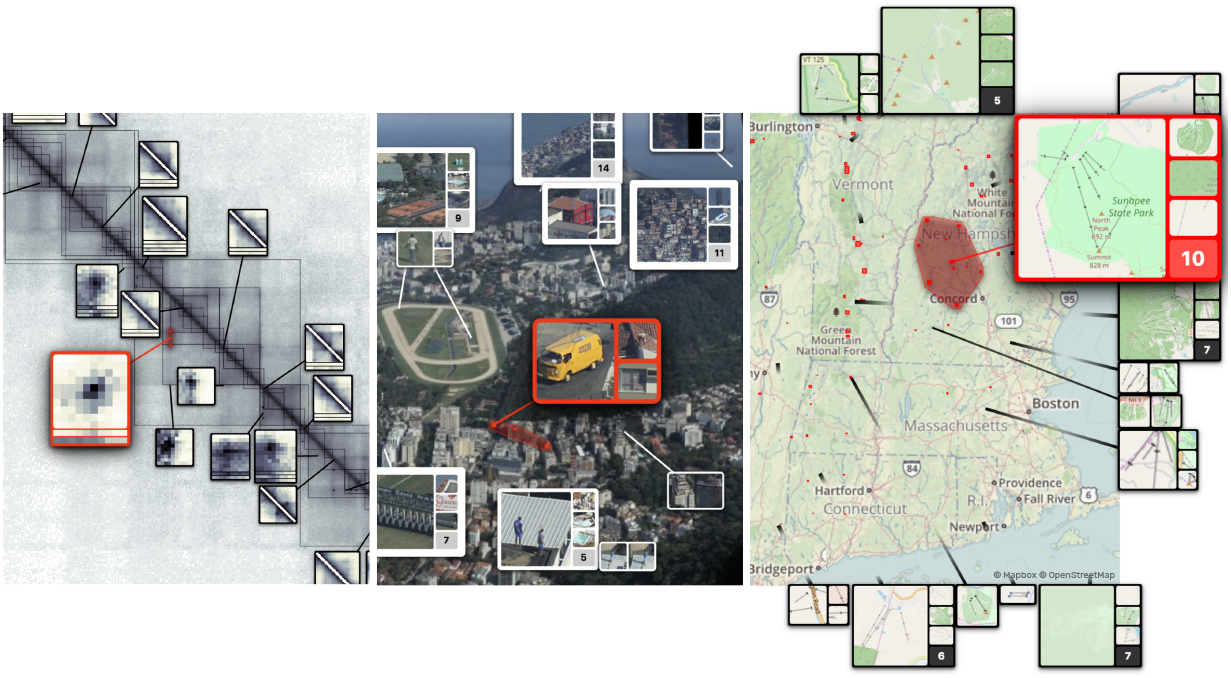
Scalable Insets applied to a genome interaction matrix [57], gigapixel image [63], and geographic map from Mapbox [43] and OpenStreetMaps [49] (left to right). Various annotated patterns are highlighted in red.

Exploring annotated patterns in their context is often needed to assess their relevance and to dissect important from unimportant regions in the visualization. For example, computational biologists study thousands of small patterns in large genome interaction matrices [12] to understand which physical interactions between regions on the genome are the driving factor that defines the 3D structure of the genome. In astronomy, researchers are exploring and comparing multiple heterogeneous galaxies and stars with super high-resolution imagery [53]. In either case, inspecting every potentially important region in detail is simply not feasible.

Exploring visual details of these annotated patterns in multiscale visualizations requires a tradeoff between several conflicting criteria. First, annotated patterns must be visible for inspection and comparison (Detail). Second, enough of the overview needs to be visible to provide context for the patterns (Context). And, third, any detailed representation of an annotated pattern should be close to its actual position in the overview (Locality). Current interactive navigation and visualization approaches, such as focus+context, overview+detail, or general highlighting techniques (section 2), address some but not all of these criteria and become difficult as repeated viewport changes, multiple manual lenses, or separate views at different zoom levels are required, which stress the user’s mental capacities.

In this paper, we describe *Scalable Insets*—a scalable visualization and interaction technique for interactively exploring and navigating large numbers of annotated patterns in multiscale visual spaces. Scalable Insets support users in early exploration through multi-focus guidance by dynamically placing magnified thumbnails of annotated patterns as insets within the viewport (Figure 1). The design of Scalable Insets is informed by interviews with genomics experts, who are engaged in exploring thousands of patterns in genome interaction matrices. To keep the number of insets stable as users navigate, we developed a technique for interactive placement of insets within the viewport and dynamic clustering of insets based on their location, type, and viewport (subsection 4.3). The degree of clustering constitutes a tradeoff between Context and Detail. Groups of patterns are visually represented as a single aggregated inset to accommodate for Detail. Details of aggregated patterns are gradually resolved as the user zooms into certain regions. We also present two dynamic mechanisms (subsection 4.2) for placing insets either within the overview (Figure 1 left and center) or on the overview’s boundary (Figure 1 right) to allow flexible adjustment to Locality. With Scalable Insets, the user can rapidly search, compare, and contextualize large pattern spaces in multiscale visualizations.

We implemented Scalable Insets as an extension to HiGlass [36], a flexible web application for viewing large tile-based datasets. The implementation currently supports gigapixel images, geographic maps, and genome interaction matrices. We present two usage scenarios for gigapixel images and geographic maps to demonstrate the functionality of Scalable Insets. Feedback from a qualitative user study with six computational biologists who explored genome interaction matrices using Scalable Insets shows that our technique is easy to learn and effective in analytic data exploration. Scalable Insets simplify the interpretation of computational results through identification and comparison of patterns in context and across zoom levels. Results from a controlled user study comparing both placement mechanisms of Scalable Insets to a standard highlighting technique provide initial evidence that Scalable Insets can reduce the time to find annotated patterns at identical accuracy and improve the accuracy in comparing pattern types. This comes at the cost of slightly slower frequency estimation of the annotated patterns. Whether this overhead is acceptable depends on the importance of the visual details of patterns for navigation.

To our knowledge, Scalable Insets is the first inset-based technique that supports the exploration of multiscale visualization where the number of insets far exceeds the available screen space and that provides first insights when an inset-based technique can provide useful and efficient guidance at scale.

**Figure 2:**
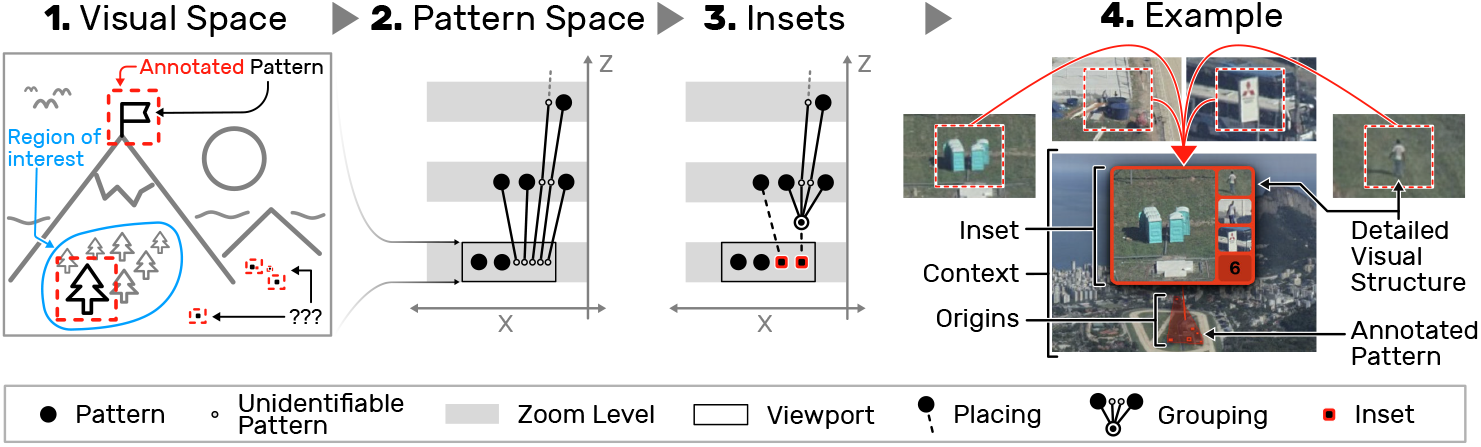
The core idea of the Scalable Insets. (1) A multiscale visualization with annotated patterns, some of which are too small to be identifiable (indicated by “???”). (2) A space-scale diagram of the virtual pattern space, showing the pattern identifiability by zoom level. (3) To provide guidance, small patterns are placed as insets into the current viewport. Scalability is ensured by dynamically grouping insets in close proximity and representing them as an aggregate as shown in (4).

## 2 Related Work

### Pan and Zoom

Pan and zoom [22] is a common technique for navigating large multiscale visualizations. Despite its widespread use, it can require a large amount of mental effort as either context or details are lost and navigating to distant regions can be time consuming [21]. Techniques have been developed to leverage the data structure to facilitate navigation and exploration. For networks, Bring&Go [48] implements navigation along a network’s links and provides navigational cues through direct visualization of a node’s neighborhood. Similarly, Dynamic Insets [23] utilizes a network topology to dynamically place off-screen nodes (i.e., annotated patterns) as visual insets inside the boundaries of the actual view. However, many data sets do not provide semantic structures to support this kind of navigation.

### Highlighting

Highlighting details is used to alleviate the lack of navigational cues [33]. For reviews on general highlighting techniques see [26,27, 39,58,66]. Irrespective of the navigation technique, knowing details about the outcome of a navigation upfront can avoid spending time on unnecessary user interactions. For instance, Scented Widgets [71] embed visual cues and simple visualizations into user interface elements. Ip and Varshney [30] describe a salience-based technique for guiding users in gigapixel images. They utilize color to highlight regions of high salience. While this is very effective to provide visual cues, these cues do not enable the user to get an understanding of the detailed visual structure of the highlighted regions without having to manipulate the viewport. Scalable Insets draws on these ideas to display the details of annotated patterns across scales.

### Aggregation and Simplification

Scalability for large visualizations can be achieved through aggregation and simplification of sets of elements. For instance, ZAME [17] is a matrix visualization technique that presents a visual summary of multiple cells at a higher level. Such a summary is composed of an embedded visualization showing the distribution of aggregated cell values. Milo et al. [47] describe network motifs, which are repetitive network patterns, to facilitate a more concise view of large networks through visual simplification [14]. Van den Elzen and van Wijk [67] integrate the ideas of ZAME and Network Simplification into a very concise overview of large networks where nodes are aggregated into small statistical summary diagrams. These techniques work well to gain an overview but the visual details of patterns are lost, which applies to many semantic zoom interfaces.

### Overview+Detail

To address the problem of lost context or missing details, one can juxtapose multiple panels at different zoom levels. The separation between these panels provides flexibility but comes at the cost of divided user attention [45]. For example, in PolyZoom [32] different zoom levels are organized hierarchically and appear as separate panels, which limits the number of regions that the user can simultaneously focus on. TrailMap [73] shows thumbnails of previously visited locations in a map visualization to support rapid bookmarking and time traveling through the exploration history. Hereby, separate panels work well as the user has already seen each location in detail before it appears as a thumbnail, but it is not clear how effective such an approach would be for guidance to new locations. HiPiler [41] supports exploration and comparisons of many patterns through interactive small multiples in a separated view. While this approach works well for out-of-context comparison, it has not been designed for guided navigation within the viewport of a multiscale visualization.

### Focus+context

Focus+context techniques show details in-place, while maintaining a continuous relationship to the context, often via distortion. The most common type of these techniques are lenses that can be moved and parameterized by the user (see [65] for a comprehensive review). For example, Table Lens [56] utilizes a degree-of-interest approach to enlarge or shrink cells of a spreadsheet. Similarly, Rubber Sheet [59] and JellyLens [54] are distortion-based techniques that enlarge areas of interest in a visualization. Melange [16] takes a different approach by folding unfocused space into a 3rd dimension like paper. Unlike our method, these techniques benefit from maintaining a continuous view at the expense of distorting the direct context around a focus region and limiting the ability to simultaneous focus on many regions.

### Detail-In-Context

Hybrid approaches magnify annotated patterns and place them as insets within the visualization [9]. For example, Pad [50] and Pad++ [3] were one of the first tools to visualize details through insets. Detail Lenses [7,34] emphasize several regions using insets that are arranged along the inner border of a map visualization. The insets are only loosely connected to their origin through sequential ordering and sparse leader lines, which ensures that the center of the map visualization is not occluded. This technique works well for up to a dozen insets but doesn’t support dynamic exploration. VAICo [60] uses insets for integrative comparison of visual details between multiple aligned images but does not concern about navigation and scalability. DragMag [70] extended the concept of Pad into a hybrid approach where magnified insets can be manually placed either within the image or on the outside border. our new technique builds on the idea of DragMag [70] and extends it with a dynamic aggregation mechanism as well as interactive exploration (Detail and Context). This allows Scalable Insets to work for cases where annotated patterns are placed very close to each other and would otherwise result in clutter.

### Label Placement

Placing insets onto an existing visualization can be compared to placing labels, e.g., on maps [72]. Many papers focus on static and point-feature label placement, i.e., the static placement of a label directly adjacent to a data point. Instead, our tool requires dynamic, excentric [20] (i.e., labels are distant from their targets), and boundary [6, 38] placement as insets are to be positioned interactively and out-of-place to avoid occluding other essential details in the visualization. Dynamic labeling [51,52,55], which has been formalized into the “consistent dynamic map labeling” and “active range optimization” problem by Been et al. [4,5], describes label placement in the context of pan and zoom interfaces, where the position and size of labels are not fixed. In order to provide real-time placement, dynamic map labeling techniques require dedicated preprocessing, which is time-consuming and, thus, hinders dynamic view composition. More importantly, Been et al. state in requirement D1 of the dynamic map labeling problem that “except for sliding in or out of the view area, labels should not vanish when zooming in or appear when zooming out” [4]. In contrast, we require that the number of insets decreases upon zooming in once the annotated patterns are large enough to be identifiable. Therefore, we’re studying an *inverted* version of the dynamic map labeling problem where insets should only be shown for annotated patterns that are too small to be identifiable to provide navigational cues and alleviate the desert fog challenge [33]. In order to address these challenges, Scalable Insets employ simulated annealing [46] as it is a generic approach [15] and can easily incorporate the requirements for Detail, Context, and Locality. Also, it has the potential to produce high-quality label placements [10] and is fast enough for interactive navigation [69].

## 3 Scalable Insets: Overview

The design of Scalable Insets is driven by the three functional requirements for detail-in-context methods. It provides a dynamic tradeoff for Detail, Context and Locality to overcome the issues one would run into with naive approaches (Figure 3). In the following, we give an overview of the technique and demonstrate the visualization and guidance aspects. Technical details on how we achieve this tradeoff are given in section 4.

**Figure 3:**
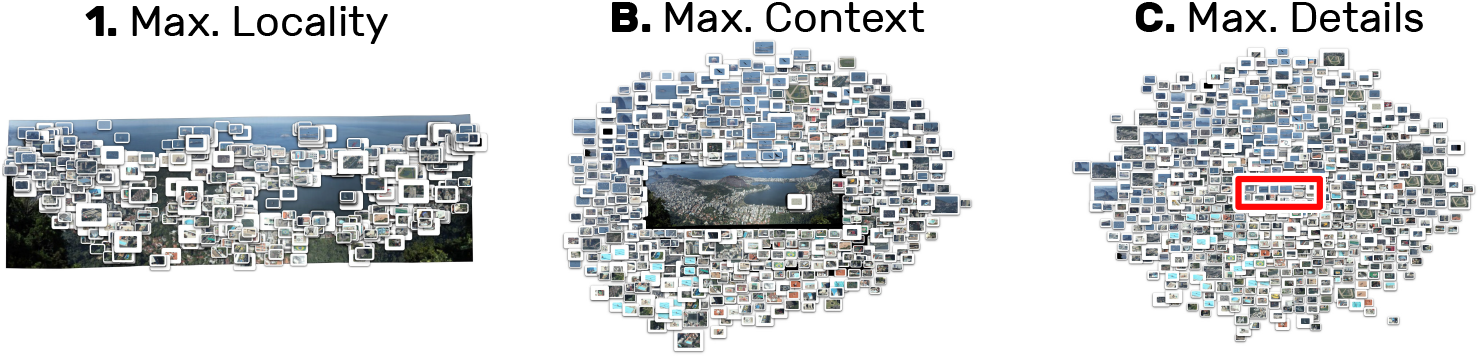
Three approaches exemplifying naive optimization of (1) Locality, (2) Context, and (3) Detail only. The red rectangle in (C) indicates the size of the occluded image for reference.

In multiscale visualizations (Figure 2.1) that contain several *annotated patterns* (henceforth just called *patterns)*, not all patterns will be identifiable at every zoom level. To this end, the notion of identifiability can be described as patterns being fully contained in the viewport and having a minimal output resolution, e.g., 24 × 24 pixels. Whenever a pattern is identifiable we are able to perceive its *detailed visual structure* (Figure 2.4). This setup lets us imagine a virtual pattern space (Figure 2.2), which defines when a pattern is visible or how much zooming is needed to identify its detailed visual structure. To reduce the interaction and navigation time to assess these visual details of patterns, we extract thumbnails of unidentifiable patterns at the zoom level that renders the pattern identifiable and place these thumbnails within the viewport as insets. The number of displayed insets can be limited by clustering several closely-located patterns into a group (Figure 2.3) and representing this group as a single aggregated inset (Figure 2.4). Together with a dynamic placement strategy that avoids occlusion of insets and patterns, this enables Scalable Insets to provide guidance for high numbers of patterns while reducing navigation time.

### 3.1 Usage Scenarios

The following two usage scenarios, on a gigapixel image and geographic map application, depict typical exploration tasks, such as searching, comparing, and contextualizing patterns, and focus likewise on overview and detail. A third, more complex use case in genomics is presented in subsection 6.2.

#### Exploring Gigapixel Images

In a gigapixel image of Rio de Janeiro [63, 64] with a resolution of 454330 × 149278 pixels (Figure 4.1) users have annotated 924 patterns such as birds, people, or cars. Some of these patterns are close together, e.g., on the same street, while others are isolated in the sea. However, most of them are not identifiable without considerate pan and zoom. We follow a hypothetical journalist who is writing an article about unseen aspects of Rio de Janeiro, which requires finding, comparing, and localizing the annotated patterns to assess “Which neighborhoods are particularly interesting to viewers?”, “What kind of patterns are most frequently annotated?”, and “Are there any unexpected patterns given their location?”.

**Figure 4:**
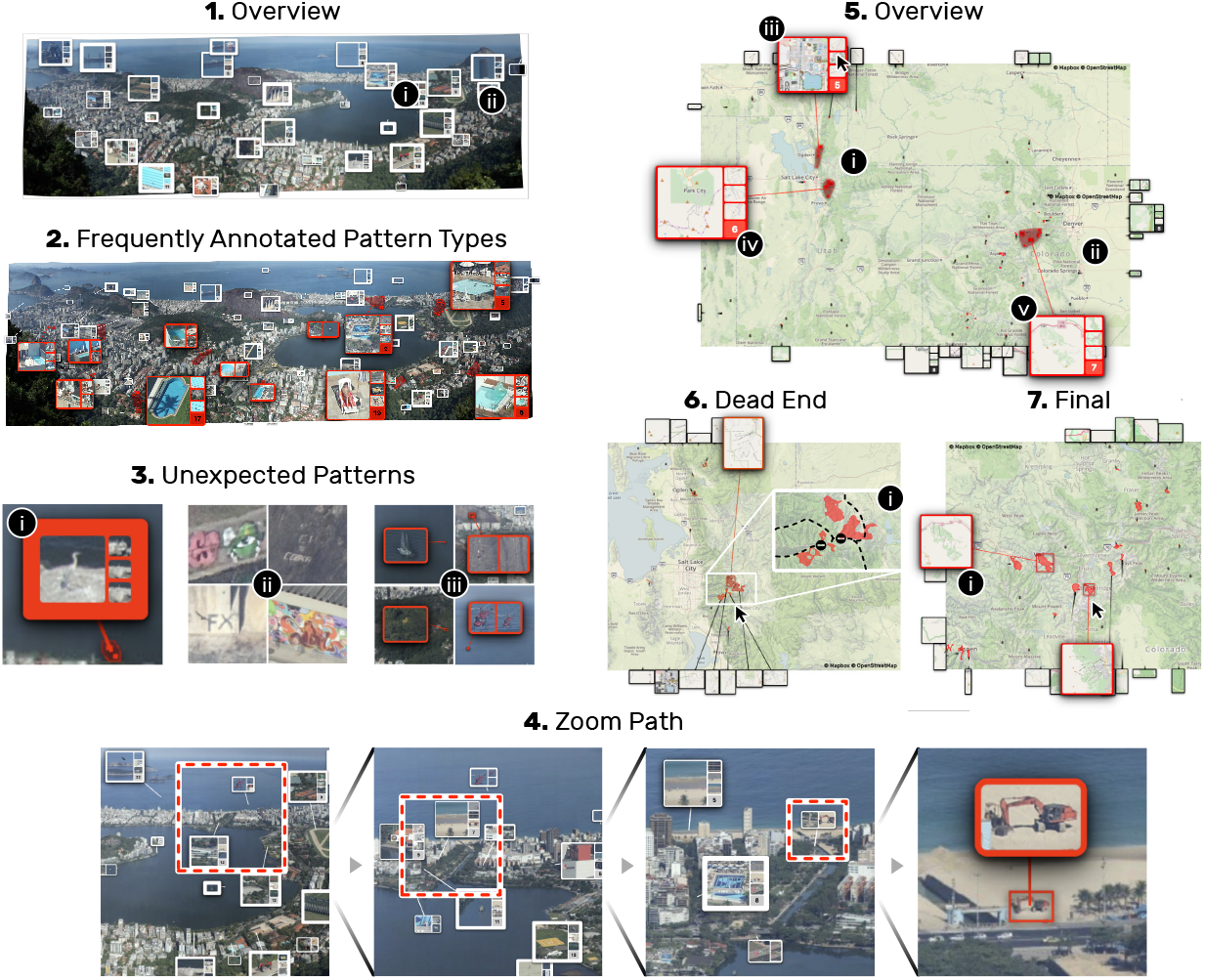
Demonstration of the Scalable Insets approach on a gigapixel image of Rio de Janeiro [63] by *The Rio—Hong Kong Connection* [64] and ski areas around Utah and Colorado shown on a map from Mapbox [43]. The screenshots illustrate how Scalable Insets enables pattern-driven exploration and navigation at scale; details are explained in subsection 3.1. See Supplementary Figures S1 and S2 for scaled-up screenshots.

Scalable Insets places insets for patterns too small to be identifiable within the viewport (Figure 4.1). Insets that would be in too close proximity to each other are grouped and represented as aggregated insets. In this example, we kept the number of insets between 25-50, which provides a good tradeoff between the Detail and Context criteria. The size of the insets ranges from 32 to 64 pixels (for the longer side) and depends on the original size of the annotated pattern, i.e., the inset that is related to the smallest pattern is 32 pixels long or wide and the inset that is related to the largest pattern is at most 64 pixels long or wide. The popularity of patterns (given by view statistics) is mapped onto the border such that thicker borders indicate higher popularity.

The journalist starts by examining the entire picture to gain an overview. At first glance, Scalable Insets reveals a relatively equal distribution of patterns, with higher frequency in areas of man-made constructions (Figure 4.1). The journalist immediately finds popular patterns as they are highlighted by a thicker border. They notice a relatively high frequency of annotated swimming pool areas (insets with a red border in Figure 4.2), which are scattered across the entire image. Upon hovering over an inset, its origin is highlighted through small red dots and a red hull in the overview. Via a click, insets are scaled up and more details of the patterns are seen. This enables the journalist to quickly identify a bird (Figure 4.3i) sitting on a locally known rock and to find interesting street art (Figure 4.3ii). Also, with Scalable Insets, several patterns located in monotone regions (Figure 4.3iii), such as the sea, are explorable with minimal pan and zoom. As the journalist inspects a specific region more closely, aggregated insets gradually split into smaller and smaller groups (Figure 4.4), presenting more details while maintaining a relatively fixed number of insets.

#### Exploring Geographic Maps

Our second scenario involves the exploration of ski areas in a geographic map from Mapbox [43] and OpenStreetMaps [49]. We obtain an estimation of the location of the world’s ski areas by analyzing *aerialway* annotations [35] from OpenStreetMap as these annotations describe potential paths of ski lifts, slopes, and other aerial ways.

The user sets out to find, compare, and localize *interesting* ski areas. Interest is defined by the size of individual ski areas and the size of multiple ski areas within close proximity. Localization of ski areas is important to determine the accessibility and proximity by car between multiple closely-located ski areas. This time, insets are shown *outside* the map to provide full access of information in the map, such as streets, cities, mountains and other important geographical information needed for localization.

The user starts exploring around Utah and Colorado (Figure 4.5). The map shows two regions with several closely-located ski areas nearby Salt Lake City (Figure 4.5i) and Denver (Figure 4.5ii). Upon scaling up an inset, the user can explore the size and shape of up to four representative ski areas among a group. This allows for fast comparison of the ski areas without the need to navigate. For example, the user compares three promising groups of several ski areas (Figure 4.5iii, 5iv, and 5v). Through hovering over different images in an aggregated inset, shown in Figure 4.5iii, the user identifies that this group contains only small ski areas as well as an outlier, i.e., a pattern that does not correspond to a ski area. While the second group (Figure 4.5iv) indeed represents several large ski areas, close inspection (Figure 4.6) reveals that the road connecting these ski areas is closed during winter (Figure 4.6i). Zooming out again, the user finds a suitable region with several larger ski areas (Figure 4.7) that are conveniently accessible by car through the Interstate 70 nearby Vail (Figure 4.7i).

**Figure 5:**
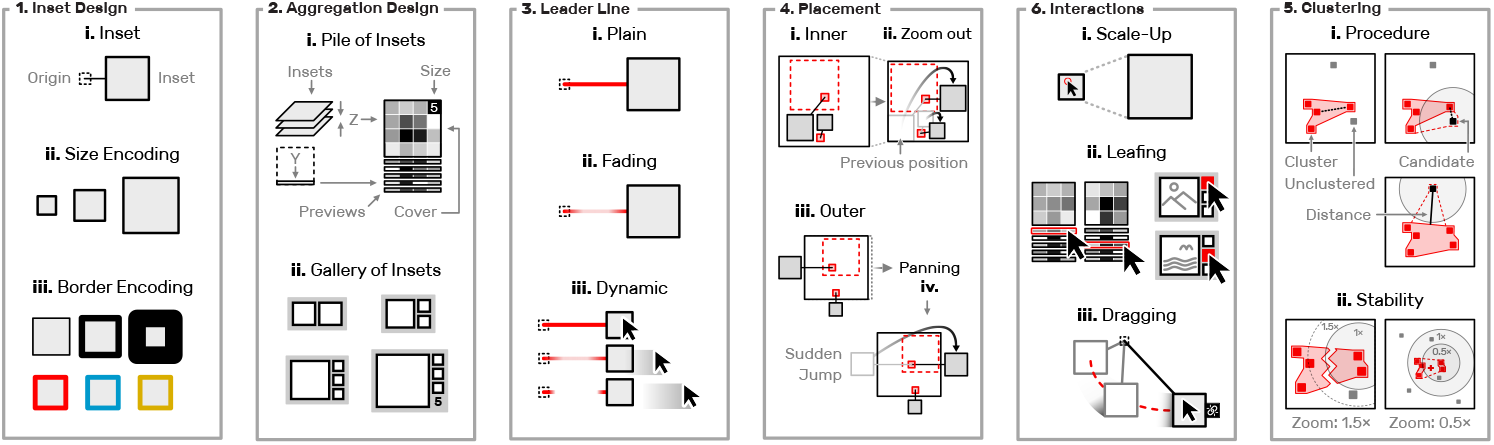
Schematic design principals of Scalable Insets. (1) Inset design and information encoding. (2) Visual representation of aggregated insets. (3) Leader line styles. (4) The inset placement mechanism and stability considerations of Scalable Insets. (5) Aggregation procedure and stability considerations. (6) Interaction between insets applied in Scalable Insets.

## 4 Scalable Insets: Technique

### 4.1 Inset and Leader Line Design

Insets *I* are small, rectangular thumbnails at an increased magnification of a subset *S* of annotated patterns *A* that are too small to be identifiable. The level of magnification is defined by the display size (in pixels) of the insets and zoom level. The display size of insets can depend on a continuous value, e.g., a confidence score or range between a user-defined minimum and maximum, but is invariant to the zoom level to give more control over Detail and Context. This comes at the cost of reduced awareness of the *depth* of annotated patterns, which we consider less important for Scalable Insets as it does not directly support finding, comparing, and contextualizing patterns. The thickness of the border can be used to encode additional information and the hue of the border is adjustable to reflect different categories of annotated patterns. Both encodings are illustrated in Figure 5.1ii and Figure 5.1iii.

A leader line is drawn between an inset *i ∈ I* and its source annotation *s_i_* ∈ *S* in order to establish a mental connection as their positions may not coincide (subsection 4.2). We designed three different leader line styles. Plain and fading leader lines are static and only differ in their alpha values along the line (Figure 5.3i and Figure 5.3ii). Dynamic leader lines adjust their opacity depending on the distance of the mouse cursor to an inset or a source annotation. Fading and dynamic leader lines minimize clutter in the event of leader line crossing to preserve context and have been shown to maintain a notion of connectedness [11,42]. We chose to limit Scalable Insets to straight leader lines for simplicity and performance reasons. While other techniques such as edge bundling [29] or boundary labeling [6] exist, the benefits are expected to be minimal. For the inner-placement, leader lines are usually very short since insets are positioned as close as possible to their source annotation. Barth et al. [2] have shown that straight leader lines for boundary labeling, which is similar to our outer-placement (subsection 4.2), perform comparably to or even better than more sophisticated methods.

After the border encoding and leader line style have been set by the user (see section 5), Scalable Insets will dynamically update the appearance based on the viewport. Hence, the visual encodings are relative to the currently visible insets.

### 4.2 Inset Placement

We have developed an algorithm for placing an inset *i* at a position *p_i_* either within the viewport (inner-placement) or at the boundary of the viewport (outer-placement), where *p_i_* is defined as a sequence of *k* moves 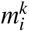 that are determined with simulated annealing [46]. The goal for moving an inset *i* is to maximize Detail and Context while minimizing Locality by optimizing a cost function *C*. This cost function depends on the current position 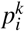 and the potential move 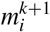 to the next position. The resulting cost determines whether a particular move from one position to another improves the placement. The final position of an inset *p_i_* is then given by a sequence of *k* moves where 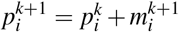 and 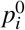 is set to the center of *i*’s source annotation.

The cost function consists of four main components. First, pairwise overlap *o_ij_* between two insets *i* and *j* should be minimized. Next, for the inner-placement pairwise overlap (*OS* and *OA*) between an inset *i*, source annotations S, and other annotated patterns *A* should be minimized. Also, insets should be placed as close to their source *s_i_* as possible, i.e., the distance *d_is_i__* between *i* and *s_i_* should be minimal. Finally, leader line crossings *l_ij_* between two insets *i* and *j* should be avoided, where *l_i_j* is 1 if the leader lines intersect and 0 otherwise. Figure 5.4i and 5.4iii illustrate both placement strategies.

Following only the above criteria could lead to drastic changes in the positioning even at minimal pan and zoom. For example, in Figure 5.4ii a subtle zoom out would lead to the occlusion of other annotated patterns (indicated as dashed, red boundaries), which could cause the inset to *jump* to the next best position. Similarly, in Figure 5.4iv subtle panning to the left would change the closest available position on the boundary and make the inset jump from one side to the other. Since these phenomena would significantly harm the usability of Scalable Insets, the Euclidean distance between an old and a new placement of *i*, denoted as *d_m_i__*, is minimized to avoid movements that would lead to only marginal improvements. In summary, the cost of moving an inset *m_i_* is comprised of the following components, where the *ws* stand for individual weights and 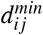 is defined as max(0, (*r_i_*+*r_j_*)−*d_ij_*) with *r_i_* being half of the diagonal of *i* to ensures a minimal distance between *i* and *j*.

- Inset distance: *D* = *w_D_d_is_i__*
- Inset movement: *M* = *w_M_d_m_i__*
- Leader line crossing: 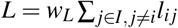
- Inset-source overlap: 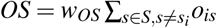
- Inset-source minimum distance: 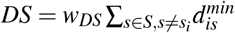
- Inset-inset overlap: 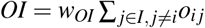
- Inset-inset minimum distance: 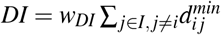
- Inset-annotation overlap: 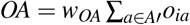
- Inset-annotation minimum distance: 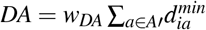

Finally, each metric is normalized to adjust for the different value ranges as follows. The distances 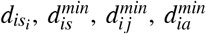 and *d_m_i__* are divided by *r_i_*. The overlap *o_ij_* between two insets is normalized by the area of the smaller inset. In contrast, the overlaps *o_is_* and *o_ia_* are normalized by the source or annotation. The intuition is that insets should ideally never overlap with other insets or their sources as this would harm Detail and Context. On the other hand, the larger an annotation is the less distracting an overlaying inset presumably is. For example, if a park in a maps visualization is annotated and spans 80% of the screen, overlaying a small 24 × 24 pixel-sized inset will presumably not harm the identifiability of the park as the inset covers only a small portion of the park. Detailed formulas and default weights are provided in Supplementary Table S1. The cost function is then defined as the sum of all components:

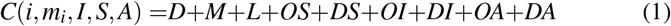

In simulated annealing the *k*-th move 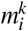 of *i* is chosen at random and accepted with a probability equal to:

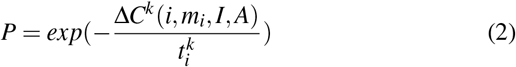

where Δ*C^k^*(*i, m_i_, I, A*) is defined as the difference between the cost of the *k*-th and the (*k*−1)-th move and 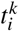 is the temperature of *i* at that move. The value of *t_i_* continuously decreases and controls the likelihood that moves, which result in higher costs, are accepted. I.e., the smaller *t_i_* the less likely it is that *i* moves to a position with higher cost. We employ exponential cooling such that 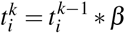, where *β* is adjustable and set to 0.8 by default. Upon zooming, the temperature of already existing insets is reset to 5% of the initial temperature to allow insets to move to better positions but avoid unnecessary moves where the cost is almost the same or worse.

Additionally, ||*m_i_*||_2_ is limited *m^max^* to avoid too large changes within one iteration of simulated annealing and decreases linearly with *t* until 0.5 * *m^max^*. Finally, in the outer-placement approach, insets are initially placed at the closest side that has the fewest insets assigned at the corresponding x or y position already. Upon pan and zoom, insets should not be moved to the opposing side as such jumps would impose high cognitive load for keeping a mental model of the insets’ positions. See Supplementary Algorithm S1 for details.

### 4.3 Aggregation

To provide scalability beyond a handful of annotated patterns and preserve Context, we have developed a density-based dynamic clustering algorithm that assigns every annotated pattern in the viewport to a particular group, called a cluster. Each cluster is represented as a single visual entity and referred to as an aggregated inset. our clustering approach is based on the spatial distance between annotated patterns in the viewport. Starting with a randomly selected pattern we find the closest cluster. only if the distance between the pattern and the bounding box of the cluster is closer than a user-defined threshold and the area of the cluster combined with the pattern is smaller than a user-defined threshold do we assign the pattern to that given cluster (Figure 5.5i). Otherwise, we create a new cluster composed of the selected pattern. See Algorithm S2 for details. Upon adding patterns to clusters, we keep a sorted list of the nearest neighbors for each newly added pattern (Algorithm S3), which will help us to determine breakpoints when zooming-in. The clustering is re-evaluated upon navigation. To improve cluster stability between short, repeated zoom changes, the distance threshold, for deciding whether an inset should be assigned to a particular cluster, is dynamically adjusted as illustrated in Figure 5.5ii. During zoom-in, clusters remain unchanged until the distance between the farthest nearest neighbors is larger than 1.5 times the distance threshold. During zoom-out, clusters and insets will not be merged until their distance is less than half the distance threshold. This limits the changes to cluster composition upon navigation to facilitate the user’s mental map of the pattern space. Details about the reevaluation algorithm are provided in Algorithm S4. our clustering approach is relatively simple to ensure high rendering performance. our approach is inspired by DBSCAN [18] but we do not implement recursive scanning of nearby neighbors as we strive for a spatially-uniform partitioning to provide useful navigational cues rather than continuous clusters.

We designed two approaches to visually represent clusters. When exploring matrices, clusters are aggregated into piles and feature a 2D cover matrix together with 1D previews [1,41]. The cover matrix represents a statistical summary of the patterns on the cluster, e.g., the arithmetic mean or variance. 1D previews are averages across the Y dimension and provide a visual cue into the variability of patterns on the pile. The number of previews is limited to a user-defined threshold. For larger numbers of previews, we employ KMeans clustering and only represent 1D previews for the average patterns of each KMeans cluster. Clusters of patterns from photographic images and geographic maps are aggregated into a gallery of cluster repre-sentatives. A small digit indicates the number of patterns for clusters larger than four. Drawing on insights from work on parameter exploration, design, or ideation space [40,44,61,62], we show a diverse set of patterns as the representatives. The largest pattern in the gallery is representing the most important pattern, according to a user-defined metric like a confidence score or click rate, or simply the area of the annotated pattern by default. The other three gallery images are chosen to represent the clusters diversity as described in detail in Algorithm S5. The pile-based aggregation is useful for patterns with well-alignable shapes, e.g., rectangles, lines, or points, to provide a concise representation of the average pattern and pattern variance. The gallery aggregation is preferable for patterns of diverse shapes as an average across shape boundaries does not provide meaningful insights.

### 4.4 Inset Interaction

Scalable Insets introduces a small number of interactions and is otherwise agnostic in terms of the navigation technique. Upon moving the mouse cursor over an inset the hue of its border and leader line change and a hull is drawn around the location of the annotated patterns represented by the inset. Upon scale-up (Figure 5.6i), it is possible to leaf through the 1D previews of a pile or the representatives of a gallery (Figure 5.6ii). Insets are draggable (Figure 5.6iii) to allow manual placement and uncover potentially occluded scenery in the overview (Context). Dragging disconnects insets from the locality criterion to avoid immediate re-positioning to the inset’s previous position upon zooming. This is visualized with a small glyph indicating a broken link (Figure 5.6iii) and can be reversed through a click on this glyph.

## 5 Implementation

To demonstrate the utility of Scalable Insets, we built a web-based prototype for gigapixel images, geographic maps, and genome interaction matrices. Scalable Insets is implemented as an extension to HiGlass [36], an open-source web-based viewer for large tile-based datasets. The Scalable Insets extension to HiGlass is implemented in JavaScript and integrates into the React [19] application framework. D3 [8] is used for data mapping and matrices are rendered in WebGL with PixiJS [25]. Scalable Insets utilizes HTML and CSS for positioning and styling insets and leader lines. Almost all parameters can be customized via a JSON configuration file^1^ and fall back to sensible default values otherwise. The server-side application, which extracts, aggregates, and serves the images for insets, is implemented in Python and built on top of Django [13]. Both, the front-end and back-end applications, are open-source, and freely available at https://github.com/flekschas/higlass-scalable-insets.

## 6 Evaluation

### 6.1 Study 1: Quantitative Evaluation

In the first user study we compare the performance of Scalable Insets to boundary boxes highlighting in terms of frequency estimation, visual search, and comparison, and we assess the effects of Context and Locality on the two placement techniques of Scalable Insets.

#### Techniques

We compared the following three techniques, which are illustrated in Figure 6. We chose bounding boxes (**BBox**) as the baseline technique given its minimal interference with the visualization, its support for areal and point annotations, and its frequent application. The method is also easy to implement and computationally simple, which enables us to evaluate the performance overhead of Scalable Insets on tasks for which the visual details of annotated patterns are irrelevant. We compare the baseline technique against Scalable Insets’ inner-(**Inside**) and outer-placement (**Outside**), where annotated patterns are shown as insets placed *inside* and *outside* the visualization respectively together with mildly-translucent boundary boxes.

**Figure 6:**
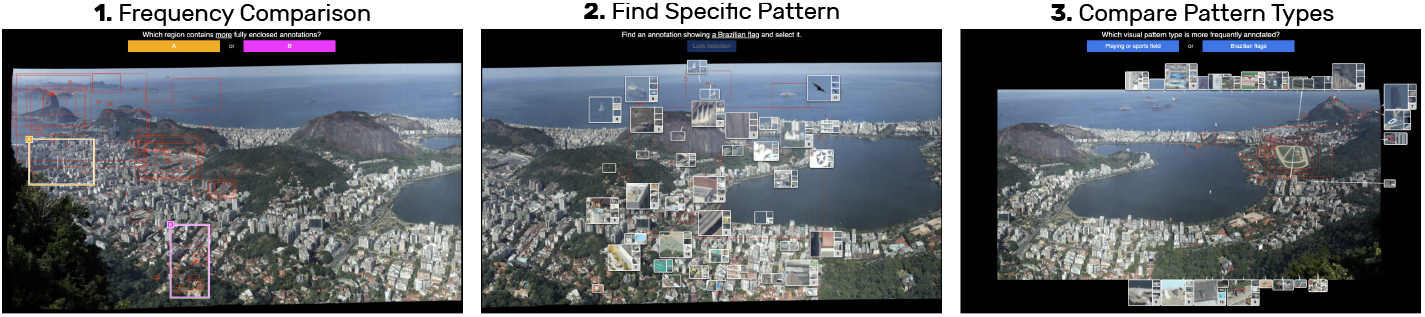
Screenshots from the first user study showing Rio de Janeiro [64] with examples of each task and technique. (1) Comparing the frequency of annotated patterns in two distinct regions with BBox. (2) Finding a specific pattern that shows a Brazilian flag with Inside. (3) Comparing the global frequency of patterns showing a “player or sports field” against “Brazilian flag” with Outside.

#### Data

For the study, we chose seven photographic gigapixel images^2^ from Gigapan [24] showing different cities (e.g., Figure 6). We used two images for practice trials and the remaining five for the test trials. The annotated patterns represent user-defined annotations from Gigapan. Image sizes ranged from 100643 × 43935 pixels to 454330 × 149278 and the number of patterns ranged from 82 to 924.

#### Tasks

We defined three tasks (illustrated in Figure 6) for which we measured completion time (in seconds) and accuracy (in percentage). In **Region** we asked the participants, *“Which region contains more fully enclosed annotations: A or B?”*. The goal of Region is to test whether the computational and visual overhead introduced by Scalable Insets impacts the general perfor-mance in exploring multiscale visualizations. Since Region does not require the user to know the visual details of annotated patterns, all techniques should perform equally. Frequency estimation is a common task for visual attention or highlighting [27] and helps to guide users [71]. Evaluating the general performance is essential as pattern exploration in multiscale visualizations consists of several tasks and worse performance in one of them could reduce the overall applicability. In **Pattern** the participants had to *“find an annotation showing* [pattern] *and select it”*. Where *[pattern]* was replaced with a description of a manually chosen pattern that appears 2–3 times in the image, e.g., “a helicopter landing field”, and is not identifiable at the initial zoom level, as this would have otherwise made the task trivial. Efficiently locating a target is a critical property of navigation and guidance techniques [32,68]. The question is whether showing visual details (Detail) of patterns too small to be visible at a certain zoom level is beneficial for search and if the distance between insets and their origins (Locality) influences the performance. In T**ype** we asked the participants *“which visual pattern type appears more frequently: A or B?”*, where *A* and *B* were replaced with two generic pattern types appearing multiple times but not equally often, e.g., “swimming pools” and “construction sites”. Most of these patterns were again not identifiable at the initial zoom level. Navigation often incorporates comparing multiple pattern instances [32] to decide which area to explore in more detail. Here we try at find out if Context and Locality influence the performance in pattern comparison. Finally, It should be noted that all tasks were conducted with interactive visualizations and especially Pattern and Type required panning and zooming to be solved accurately.

#### Hypotheses

We formulated one hypothesis per task: First (**H1**), there will be no significant difference in time and accuracy between any of the techniques for Region as the detailed visual structure of annotated patterns is not important for estimating the pattern frequency. Second (**H2**), for the Pattern task, Inside will be faster than Outside and Outside will be faster than BBox. We expect the inner-placement of insets to be most efficient as the detailed visual structure of annotated patterns is displayed spatially close to their original location, i.e., eye movement is minimized. We expect the outer-placement of insets to be slightly slower compared to Inside, due to eye movement, but faster than the baseline technique as it still shows the visual details of annotated patterns. Finally (**H3**), for the Type task, Inside and Osutside will be faster than BBox. We expect Scalable Insets with both placement mechanisms to perform equally fast and better than BBOX as they both highlight the detailed visual structure of annotated patterns.

#### Participants

We recruited 18 participants (7 female and 11 male) from Harvard University after obtaining approval from Harvard’s Institutional Review Board. 3 participants were aged between 18 to 25, 13 were aged between 25 to 30, and the remaining 2 were aged between 31 and 35. All participants volunteered, had no visual impairments, and were not required to have particular computer skills. Each participant received a $15 gift cardupon completion of the experiment.

#### Study Design

We used the following full-factorial within-subject study design with Latin square-randomized order of the techniques:

|  |  |  |
| --- | --- | --- |
|  | 18 | participants |
| × | 3 | techniques: BBOX, INSIDE, OUTSIDE |
| × | 3 | tasks: REGION, PATTERN, TYPE |
| × | 5 | timed repetition (excluding 2 practice trials) |
|  | 810 | total trials (45 per participant) |

Participants were split evenly between the 3 Latin square-randomized technique orders. The task order was kept constant across all participants and conditions. To avoid learning effects between images, the set of annotated patterns was split into 3 groups of an equally-sized region in the image. The order of these regions was kept constant, i.e., the first technique always operated on the first quadrant of the images. To avoid memory effects between Region and Pattern, we excluded the patterns relevant in Pattern from the Region trials. Furthermore, the patterns for Pattern were chosen such that they are not contained in or in close proximity to the two regions that were compared in REGIoN. Each trial is repeated on the 5 different images. Finally, we ordered the images by size and amount of patterns. The first image is the smallest and contains the fewest annotations. The annotation frequency, size, and structural difficulty increase with the last image being the largest and most frequently annotated. The order of the images was kept the same.

#### Setup

The study was conducted on a MacBook Pro (Table S7), which was equipped with a standard two-button scroll-wheel mouse. Inside and Outside parameter settings are provided in Table S5.

#### Procedure

The study was conducted in individual sessions that started with an overview of the general procedure, obtaining consent, and briefly introducing the study software (2 minutes), which guided the participants through each task. In the beginning, gender and age group was collected and participants had to solve a 12-image Ishihara color blindness test [31]. Prior to the actual test trials, participants were shown detailed instructions and had to complete two practice trials to familiarize themselves with the user interface and respective task. once the user selected an answer the timer stopped and a click on the next button was awaited before starting the next trial. Participants were instructed to finish the trials *as fast as and as accurately as possible* but to also *rely on their intuition* when estimating frequencies (Region and Type). In the end, participants were asked to anonymously fill out a questionnaire on the general impression of Scalable Insets.

#### Results

We found that completion time was not normally distributed after testing goodness-of-fit of a fitted normal distribution using a Shapiro-Wilk test. We visually inspected dot plots with individual data points and removed trials that are more than 4 standard deviations away from the arithmetic mean time. These trials are most likely related to severe distraction. This resulted in the removal of 3 trials from Region, 1 trial from Pattern, and 1 trial from TYPE. Given the non-normal distribution of completion time and the unequal number of trials due to outlier removal, we report on non-parametric Kruskal-Wallis and post-hoc Holm–Bonferroni-corrected [28] Mann-Whitney U tests with for assessing significance. For accuracy, we used a Chi-square test of independence. All *p*-values are reported for *α* = .05. In the following, we report on time (in seconds) and accuracy (in percent) by task and use pp to denote percentage points. The results are summarized in Figure 7.

**Figure 7:**
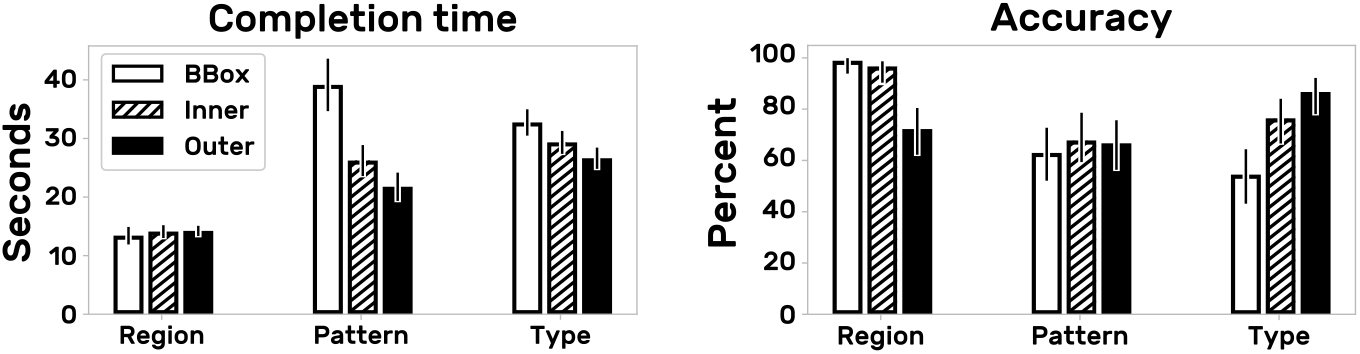
Mean completion time in seconds (lower is better) and mean accuracy in percent (higher is better) across tasks and techniques. Error bars indicate the standard error. Note, due to non-normal distribution of completion time we report significance on the median time using Kruskal-Wallis and Holm-Bonferroni-corrected Mann-Whitney U tests.

#### Results for Region

A pairwise post-hoc analysis revealed a significant difference for completion time between BBox-Inside (p=.0187) and BBox-Outside (*p*=.0031) but not for Inside-Outside. The respective mean times are BBox=13.4s (SD=14.2), Inside=14.1s (SD=10.4), Out-side=14.2s (SD=8.25). This constitutes an approximate speedup of 5.0% for BBox over Inside and 5.6% for BBox over Outside. These results let us reject H1 as BBox was fastest. Given that in absolute numbers Inside and Outside are only 0.7s and 0.8s slower than BBox suggests that overhead imposed by insets is fairly small and likely diminishes upon performance improvements to the current implementation of Scalable Insets as discussed later. For accuracy, we found a significant increase for BBox (27pp) and Inside (24pp) over Outside (*χ*^2^(1,N=177)=30.4, *p* <.0001 and *χ*^2^(1,N=179)=22.7, *p* <.0001). We believe that this difference might be caused by misinterpretation or confusion of leader line stubs as they might have appeared like or occluded the bounding box around annotated patterns.

#### Results for Pattern

We found marginally significant differences for completion time between BBox-Inside (*p*=.057) and significant difference between BBox-Outside (*p*=.0018). The mean times were BBox=39.4 (SD=42.5), Inside=26.2 (SD=25.1), Outside=21.8 (SD=22.9). This amounts for an approximate 44.9% speedup for Outside over BBox and 33% speedup for Inside over BBox and shows high potential for search tasks when the detailed visual structure of patterns and their location is important. While we could not confirm the superiority of Inside over Outside, we find a clear improvement of Scalable Insets over the baseline technique BBox. We can thus partly accept H2. The speedup is a very strong indicator that the core principal of Scalable Insets is efficient for pattern search. We hypothesize that the stronger speedup for Outside might be due to the alignment of insets in the outer-placement mechanism as this is potentially beneficial for fast sequential scanning. This advantage might diminish when contextual cues are included in the search task as well, e.g., find an annotated car at an intersection, which we did not explicitly test for. For accuracy, we did not find any significant differences.

#### Results for Type

We found no significant differences for completion time between BBox, Inside, and Outside. The mean times were BBox=32.7 (SD=21.3), Inside=28.3 (SD=16.1), Outside=26.6 (SD=17.2). Although only marginally significant, we recognize an approximate 18.7% speedup for Outside over BBox on completion time. These results let us reject hypothesis H3. For accuracy, our results show pairwise significant differences between BBox-Inside (*χ*^2^(1,N=179)=9.66, *p*=.0022) and BBox-Outside (*χ*^2^(1,N=180)=23.5, *p* <.0001). This describes an approximate improvement of 22pp for Inside over BBox and an approximate improvement of 32pp for Outside over BBox. While the completion time alone is not conclusive, the results for accuracy indicate that Outside and Inside provide a much better understanding of the distribution of pattern types. Participants with BBOX were only marginally significantly slower but made a lot more mistakes.

Finally, the 5 repetitions were completed on different images. Since all participants performed each task with every design on all images, the differences between images have equal impact on all designs and the comparison between the 3 designs indicates true differences. Intra-task variation is not our concern but it would be an interesting research question for future work.

#### Qualitative feedback

In the closing questionnaire (Supplementary Table S2), participants were asked to rank the general impression (Q1), usefulness (Q2), and simplicity to learn (Q3) on a 5-point Likert scale ranging from strongly disagree or negative (1) to strongly agree or positive (5) (Figure 9). Overall, the participants perceived the Scalable Insets approach as promising (Q1) and useful (Q2) for exploration. The high ratings for the usability (Q3) indicate that the participants had no problem learning how to use Scalable Insets. In the free-form questions on intuition and general feedback (Q4 and Q5), two aspects were mentioned multiple times: sudden disappearance of insets once the size of the original location is larger than a pre-defined threshold (Q4) and the relatively low resolution of insets until scale-up (Q5). The first aspect could be addressed in the future by employing a doi function for dynamically adjusting the behavior of insets. The relatively low resolution was due to performance reasons and can be mitigated through preprocessing of the inset’s images.

### 6.2 Study 2: Qualitative Evaluation

The goal of our second study was to evaluate the usability and usefulness of Scalable Insets in a scientific setting. To that end we conducted exploratory sessions with computational biologists, exploring large-scale matrices from structural genomics (Figure 8) using Inside and Outside. The apparatus was the same as in the first study.

**Figure 8:**
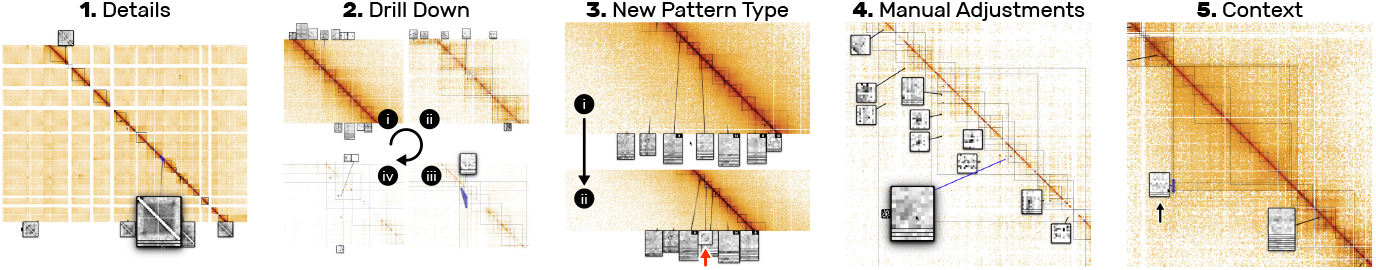
Notes from the second user study. (1) Detailed inspection of an unexpected pattern through scale-up. (2) Zoom into the original location of a clustered pile of insets until it disperses. (3) Upon zoom-out, a new pattern appeared as an inset (see red arrow) and was recognized immediately. (4) Manual inspection of the context around the pile’s origin (end of the blue line). (5) Focus on a pile of two insets due to their location.

#### Dataset

We obtained genome interaction matrices at the order of 3× 3 million rows and columns that are visualized as a heatmap. These heatmaps contain various types of visual patterns that act as proxies for cellular properties and functions and are annotated algorithmically. The frequency of these patterns ranges from a few hundred to hundreds of thousands per matrix, and analysts are interested in finding, comparing, and contextualizing patterns across many zoom levels (more details are described elsewhere [41]).

#### Participants

We recruited 6 computational biologists (2 female and 4 male) working in structural genomics. 3 experts are PhD candidates and 3 are postdoctoral researchers. Every expert was familiar with HiGlass [37] but did not see Scalable Insets. All participants volunteered, had no visual impairments, and received $15 upon completion.

#### Tasks & Study Design

The study consisted of individual open-ended sessions lasting between 20 to 30 minutes. The domain experts were asked to verbalize their thoughts and actions. Most participants started with Inside and some with Outside. In both cases the participants switched the layout after half the time.

#### Procedure

After collecting consent, each participant was briefly introduced to Scalable Insets and the data that was to be explored (<2 minutes). Next, we asked the participants to freely explore the data while the screen and audio was recorded for later analysis. Finally, each participant anonymously filled out a questionnaire.

#### Results

The results suggest that Scalable Insets is easy-to-learn as the participants immediately picked up the core concept of Scalable Insets and started exploring the dataset. Having magnified and aggregated views of annotated patterns Inside the visualization was perceived useful for exploring genome interaction matrices. The domain experts noted that they were able to find and evaluate the detailed visual structure of the patterns without having to navigate extensively. See Supplementary Table S3 for the complete protocol.

In general, we found that insets were often used as quick previews to assess a pattern before zooming into a specific location. Frequently, this assessment included comparison between different patterns and involved sequential scale-ups of the compared insets via mouse clicks. Also, some participants first sequentially hovered over the insets or moved the mouse cursor along the diagonal of the matrix to localize the insets. During pan-and-zoom, we noticed that participants used insets as navigational guidance by hovering the *target* inset, which subsequently highlighted the original location of the annotated pattern. Furthermore, we observed that Inside and Outside led to different behavior. For Outside, participants zoomed less often into the original location of an inset and instead compared the visual details of the patterns more frequently. P3 noticed, *“It’s easier to compare [the patterns] since they are all lined up nicely.”*. Some participants preferred one placement approach over the other but everyone noted that it would be useful to have the ability to switch between placement modes interactively. Some participants needed time to get used to the Outside with fading leader lines but appreciated the benefit for context preservation quickly. A drawback of the current implementation, mentioned by many participants, is the sudden disappearance of insets once their original location is large enough (e.g., 24 × 24 pixels). Although it is necessary to remove insets to release visual space for other annotated patterns, it could be beneficial to adjust the threshold based on a doi function. Insets in close proximity to the mouse cursor could remain visible until the user changes its focus.

Finally, as in the first user study, participants rated (Figure 9) how useful and easy it was to learn Scalable Insets (Table S4 Q1-3) and also compared the functionality and usefulness for the domain-specific application (Table S4 Q5-10). The domain experts strongly indicated that they would use Scalable Insets to explore their own data, given that additional application-specific features are added. For example, P4 would like to “pin” insets to compare and aggregate them with other far-away annotated patterns. Others asked for the ability to dynamically change the color map, resolution, or size of an inset as well as annotating patterns interactively.

**Figure 9:**
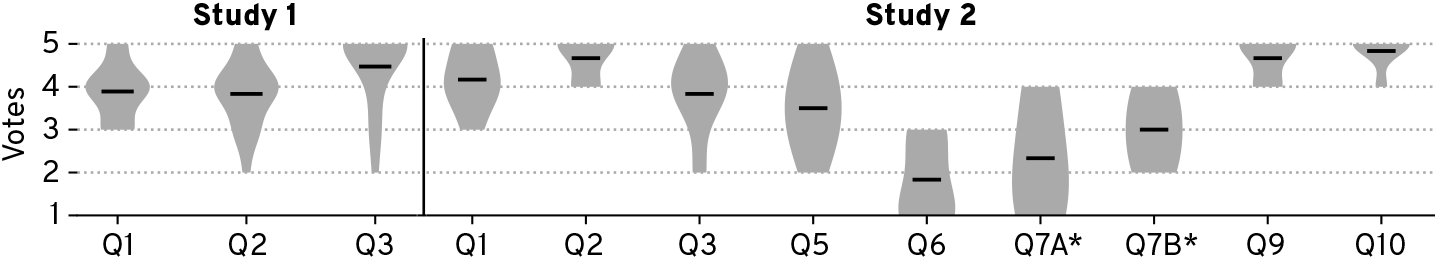
Results of the closing questionnaire. Mean values are indicated by a black bar. Questions marked with an asterisk have low absolute votes and are inconclusive. For details see Supplementary Table S6.

### 6.3 Computational Performance

We conducted a preliminary performance analysis of Scalable Insets’ placements and clustering algorithms on the gigapixel image shown in Figure 6 with all 924 annotations. We decided to focus only on the placement and clustering algorithms as those are the core contributions of this paper. The choice of preprocessing, representation sampling, aggregation, or data transfer has an additionally impact on the performance but is highly application dependent. For reproducibility, we used a scripted navigation trajectory that we interpolated with D3 [8] zoom. We ran the animation 10 times and measured the frames per second (FPS) in Google Chrome’s DevTools (v74) on the same computer from subsection 6.1. Table 1 provides a summary of the FPS across the animated trajectory that is shown in detail in Supplementary Figure S3. The inner- and outer-placement show an average frame rate of 31 and 41 FPS respectively. The frame rate drops noticeably when more than 30 insets are displayed, the overall number of annotations gets close to 1000, or the location changes markedly in a short amount of time. Without Scalable Insets the animation runs at 60 FPS.

**Table 1:** Summary of the performance analysis. Outside has a slightly higher frame rate. For details see Supplementary Figure S3.

| Time (s) | 0 | 2 | 4 | 6 | 8 | 10 | 12 | 14 | 16 | 18 |
| --- | --- | --- | --- | --- | --- | --- | --- | --- | --- | --- |
| **FPS Inner** | 14 | 15 | 35 | 51 | 32 | 21 | 47 | 45 | 23 | 24 |
| **FPS Outer** | 22 | 31 | 54 | 59 | 38 | 40 | 59 | 53 | 25 | 32 |
| **\|Insets\|** | 42 | 40 | 33 | 18 | 14 | 36 | 22 | 15 | 26 | 30 |
| **\|Annotations\|** | 924 | 893 | 492 | 123 | 120 | 281 | 361 | 116 | 165 | 674 |

## 7 Discussion

We designed Scalable Insets as a guidance technique to improve exploration and navigation of high numbers of annotated patterns for tasks that involve awareness of the patterns’ detailed visual structure, context, and location. As the first study indicates, there is strong evidence that Scalable Insets support pattern search and comparison of pattern types. The second study found that the choice of placement depends on the importance of overview and context; inner-placement was preferred for contextualizing annotated patterns, while outer-placement was preferred for pattern comparison and gaining an overview.

### Scalability and Limitations

Scalable Insets has been designed for dense but sparsely-annotated multiscale visualizations where every pixel is associated with a data point. The current prototypical implementation can simultaneously visualize and place up to 100 insets from up to 1000 annotations within a viewport. The performance can be improved in the future through advanced preprocessing or more extensive use of WebGL. In general, the usefulness and performance of Scalable Insets decrease when the clusters of aggregated insets get significantly greater than 10 over several zoom levels. Also, the total number of insets should be limited to at most 50 to avoid low FPS and high cognitive load. Therefore, applying Scalable Insets to densely-annotated visualizations would require additional features such as filtering to ensure effective guidance.

### Tradeoff Between Details, Context, and Locality

Scalable Insets set out to provide a tradeoff between Detail, Context, and Locality to manage exploration of high numbers of annotated patterns but to this end, the tradeoff is configured upfront by employing sensible defaults for the three use cases presented in this paper. To determine the parameters for placement and clustering, we manually inspected the visualization at an overview and detail viewpoint a few times to balance Detail and Context. For unevenly distributed annotated patterns we loosened the Locality requirement to make use of areas without annotations. An unsolved question beyond the Scalable Insets technique is what defines a “good” tradeoff and how could this tradeoff be adjusted interactively during navigation and exploration depending on the user’s task.

### Inset Design and Cluster Representation

The design of the insets content highly depends on the data type and specific tasks. We provide two generic approaches: piling for pattern types of homogeneous shape, such as dots or blocks in matrices, and a gallery view of cluster representatives for pattern types of high variance and diverging shapes, such as patterns found in images and geographic maps. As participants in both studies noted, there are further possibilities for application-specific cluster representations. For instance, we employ a relatively simple representative sampling technique (Algorithm S5) based on Euclidean distance for visual diversity and performance. It would be interesting to study other types of sampling techniques in future work that incorporate the semantics of the underlying pattern.

### Generalizability

Scalable Insets is not limited to the three data types presented in this paper. Our technique can be applied to any types of multiscale visualization that exhibit a large number of sparsely-distributed patterns. Even mono-scale visualizations that entail a third dimension, such as magnetic resonance imaging, could be enhanced with Scalable Insets. The effectiveness of Scalable Insets depends on how important the detailed visual structure, context, and location of the annotated patterns is. Finally, Scalable Insets is designed as a guidance technique with a minimal interaction space to be compatible with a wide range of navigation techniques like PolyZoom [32].

## 8 Conclusion and Future Work

Scalable Insets enable guided exploration and navigation of large numbers of sparsely-distributed annotated patterns 2D multiscale visualizations. Scalable Insets visualizes annotated patterns as magnified thumbnails and dynamically places and aggregates these insets based on location and pattern type. While Scalable Insets currently supports images, maps, and matrices, we plan for other data types and scenarios, investigate techniques to cope with dense regions of patterns, and support more free-form exploration, e.g., through pinning and manually grouping of insets.

## Supporting information

Screencast

## Acknowledgement

We wish to thank all participants from our two user studies who helped us evaluate Scalable Insets and provided useful feedback. This work was supported in part by the National Institutes of Health (U01 CA200059 and R00 HG007583).

## Footnotes

1 https://github.com/flekschas/higlass-scalable-insets#config

2 Gigapan IDs: 149705, 40280, 48635, 47373, 33411, 72802, and 66020

