## Supplementary Material for "Pattern-Driven Navigation in 2D Multiscale Visualizations with Scalable Insets"

Fritz Lekschas

Harvard School of Engineering and Applied Sciences

Michael Behrisch

Harvard School of Engineering and Applied Sciences

Benjamin Bach

University of Edinburgh

Peter Kerpedjiev

Harvard Medical School

Nils Gehlenborg

Harvard Medical School

Hanspeter Pfister

Harvard School of Engineering and Applied Sciences

#### Supplementary Figures

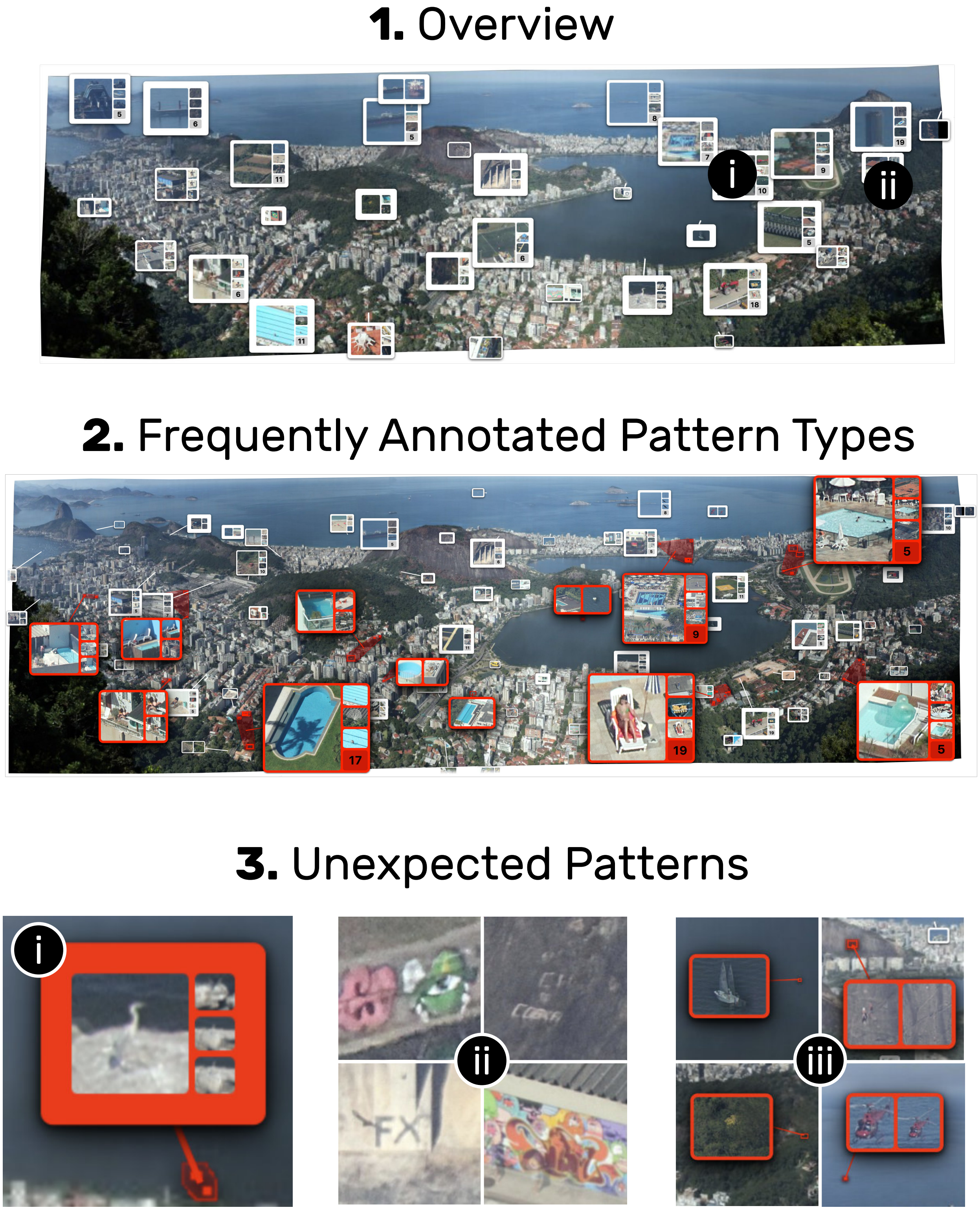


Figure S1: Scaled-up version of Figure 3.1-3

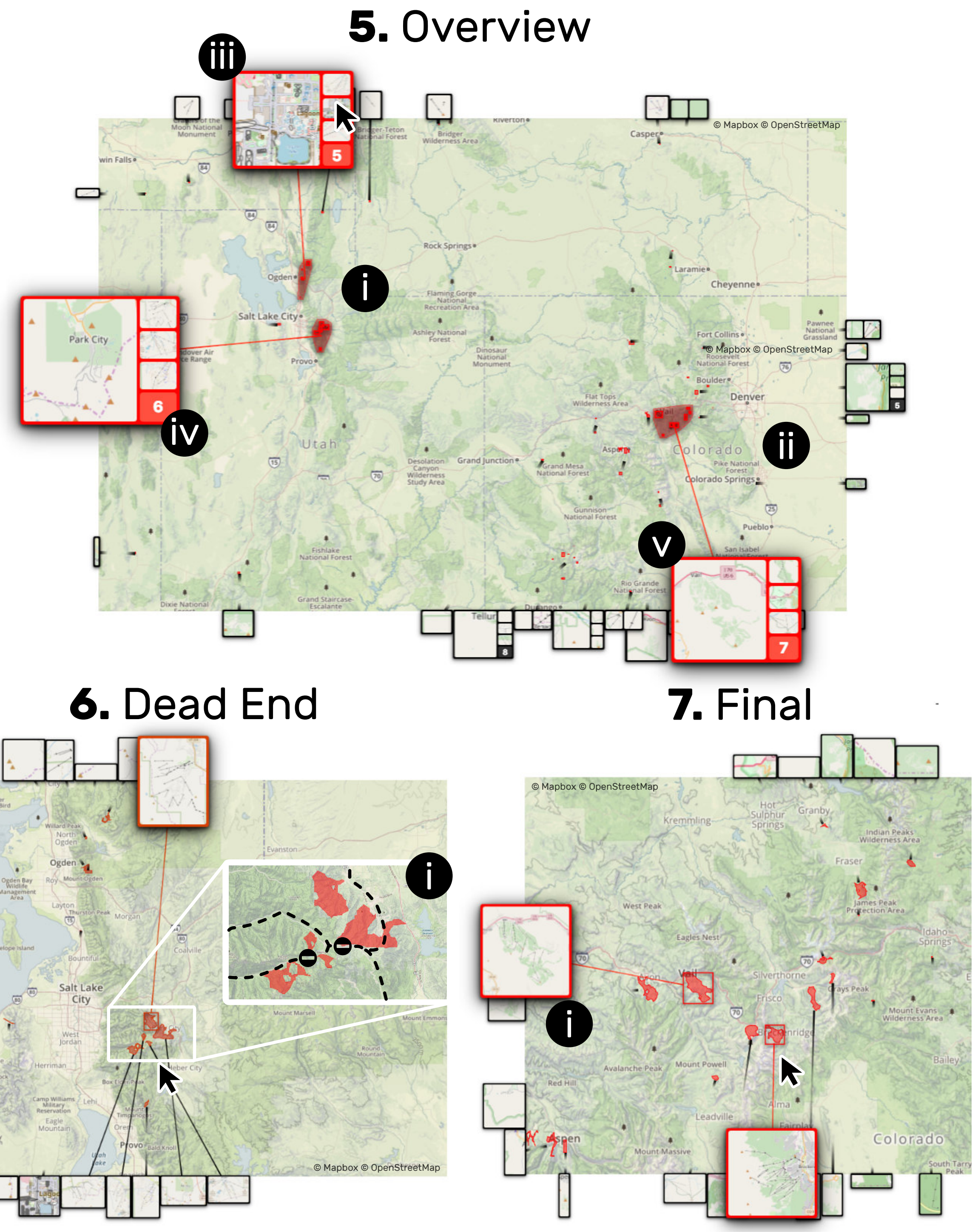


Figure S2: Scaled-up version of Figure 3.5-7

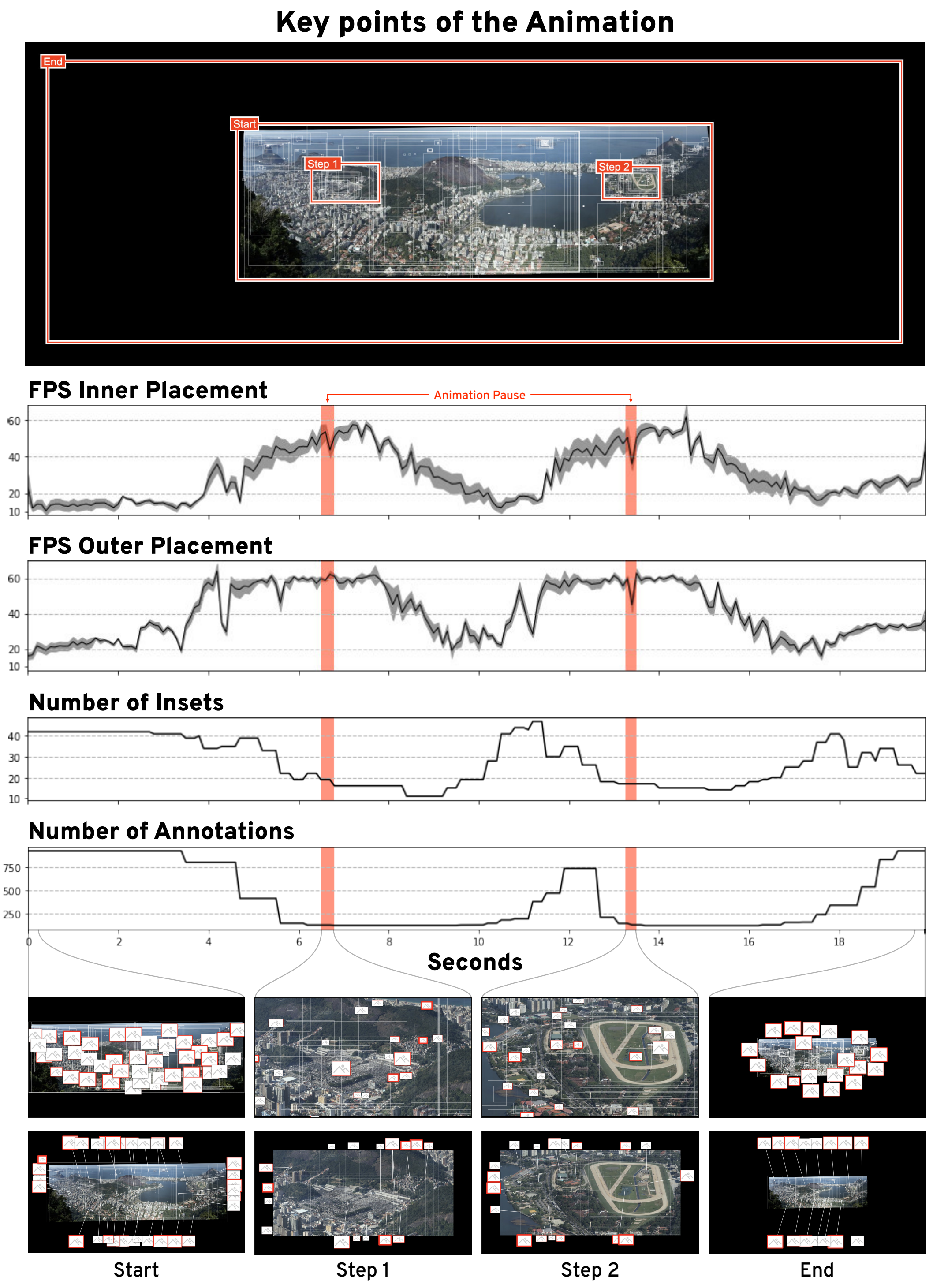


Figure S3: Detailed view of the performance analysis explained in Section 6.3. The gray area in "FPS Inner Placement" and "FPS Outer Placement" shows the standard error. The two regions highlighted in red mark 250-millisecond pauses of the animated transition between the key points. For static views Google Chrome's DevTool v74 reports very low frame rates as the frames are not refreshed. Therefore the FPS during these two pauses should be ignored.

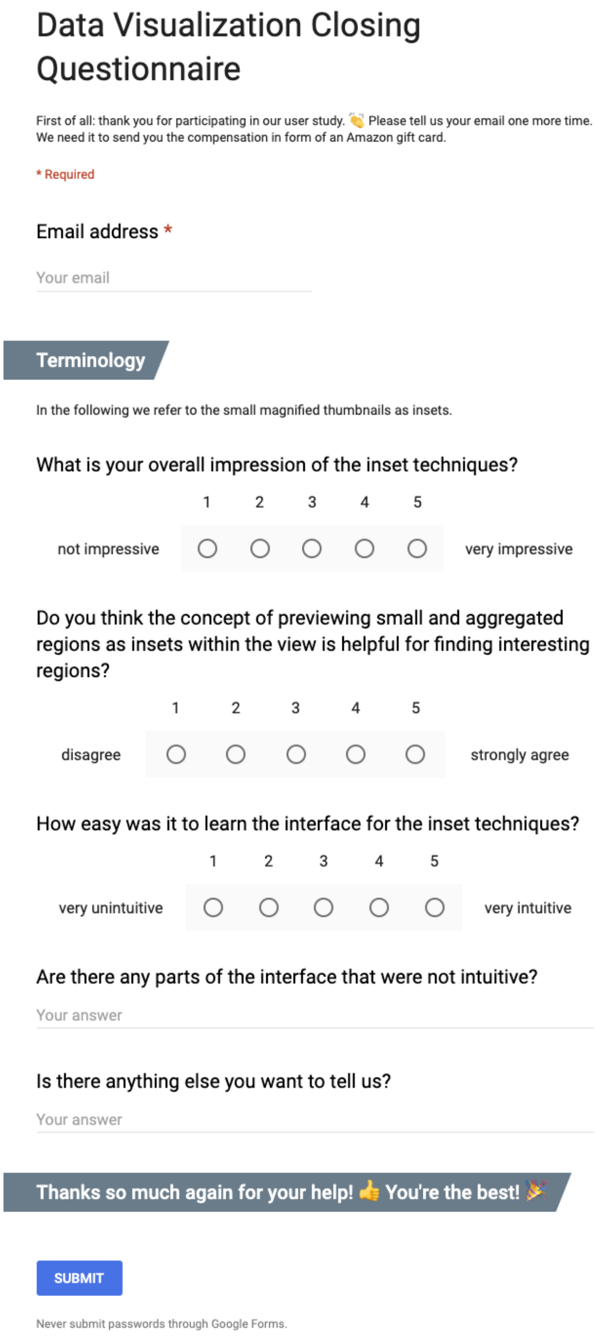

### Data Visualization Closing Questionnaire

First of all: thank you for participating in our user study. 👏 Please tell us your email one more time. We need it to send you the compensation in form of an Amazon gift card.

* Required

Email address *

Your email

#### Terminology

In the following we refer to the small magnified thumbnails as insets.

What is your overall impression of the inset techniques?

| | 1 | 2 | 3 | 4 | 5 | |
|---|---|---|---|---|---|---|
| not impressive | ○ | ○ | ○ | ○ | ○ | very impressive |

Do you think the concept of previewing small and aggregated regions as insets within the view is helpful for finding interesting regions?

| | 1 | 2 | 3 | 4 | 5 | |
|---|---|---|---|---|---|---|
| disagree | ○ | ○ | ○ | ○ | ○ | strongly agree |

How easy was it to learn the interface for the inset techniques?

| | 1 | 2 | 3 | 4 | 5 | |
|---|---|---|---|---|---|---|
| very unintuitive | ○ | ○ | ○ | ○ | ○ | very intuitive |

Are there any parts of the interface that were not intuitive?

Your answer

Is there anything else you want to tell us?

Your answer

#### Thanks so much again for your help! 👍 You're the best! 🎉

SUBMIT

Never submit passwords through Google Forms.

Figure S4: A screenshot of the closing questionnaire of user study 1 for completeness. The questionnaire was realized with Google Forms. The results are presented in Table S2.

#### Supplementary Tables

| Component | Weight | Unit | Normalized By | Formula |
|---|---|---|---|---|
| inset distance | 1.0 | pixels | inset radius | $\|\|i^c - s_i^c\|\|/r^i$ |
| inset–inset overlap | 2.0 | pixels$^2$ | inset area | $\text{overlap}(i, j)/\min(i^a, j^a)$ |
| inset–source overlap | 2.0 | pixels$^2$ | source area | $\text{overlap}(i, s_i)/s_i^a$ |
| inset–annotation overlap | 0.5 | pixels$^2$ | annotation area | $\text{overlap}(i, a)/a^a$ |
| inset–inset min. distance | 0.5 | pixels | inset radius | $\max(0, (r_i - \|\|i^c - j^c\|\|)/r_i)$ |
| inset–source min. distance | 0.5 | pixels | inset radius | $\max(0, (r_i - \|\|i^c - s_i^c\|\|)/r_i)$ |
| inset–annotation min. distance | 0.25 | pixels | inset radius | $\max(0, (r_i + b_a - \|\|i^c - a^c\|\|)/r_i)$ |
| leader line intersection | 1.0 | count | *none* | $\text{intersect}(i, j)$ |
| move distance | 0.5 | pixels | inset radius | $\|\|m_i\|\|/r^i$ |

Table S1: Components with their default weights and normalization terms. The notation follows Section 4, where $i$ and $j$ represent two insets. Additionally, $i^c$, $s^c$, and $a^c$ stand for the location of the center of an inset, an inset's source annotation, or another annotation. The area of an annotation is denoted by $s^a$ or $a^a$ and the breadth an annotation is denoted by $b_a$. We differentiate between "sources" and "annotations", where "sources" are annotations too small to be identifiable and are associated with insets and "annotations" are all other annotations that are large enough to be considered identifiable. The inset radius $r_i$ is defined as $\sqrt{(0.5 \times width)^2 + (0.5 \times height)^2}$.

| **1** | **2** | **3** | **4** | **5** |
|---|---|---|---|---|
| 1. What is your overall impression of the inset techniques? (1: not impressive; 5: very impressive) | | | | |
| | | xxxxx | xxxxxxxxxxx | xxx |
| 2. Do you think the concept of previewing small and aggregated regions as insets within the view is helpful for finding interesting regions? (1: disagree; 5: strongly agree) | | | | |
| | x | xxxx | xxxxxxxxxxx | xxx |
| 3. How easy was it to learn the interface for the inset techniques? (1: very unintuitive; 5: very intuitive) | | | | |
| | x | xx | xx | xxxxxxxxxxxxx |
| 4. Are there any parts of the interface that were not intuitive? | | | | |
| • Inset disappearance (4x)<br>• Manually show an inset through a click on the bounding box of the source annotation<br>• Pile size number is confusing as it's not always visible<br>• Make aggregated insets dispersable<br>• Foreign terminology (copula)<br>• Enlarge insets on zoom (i.e., the mouse wheel event)<br>• Inset movement was distracting (inner-placement)<br>• Insets were very prominent<br>• Difference between cluster representatives is not obvious | | | | |
| 5. Is there anything else you want to tell us? | | | | |
| • Low resolution of insets (3x)<br>• Cultural bias needs to be taken into account<br>• Scroll acceleration is too fast<br>• Make aggregated insets dispersable | | | | |

Table S2: Closing questionnaire of the first user study. Each "x" stands for the answer of one of the participants. The lists for question 4 and 5 have been paraphrased and combined for brevity. The questionnaire was realized with Google Forms and translated into this table. For completeness please see a screenshot of the Google Form in Figure S4. The wording and values are identical. We employed end labeling, i.e., only points 1 and 5 were labeled as shown above. All questions were optional.

| Participant | Action |
|---|---|
| P1 | *Started with inner-placement* |
| | • Slow panning to gain an understanding of the view and data |
| | • Gently zoomed to an inset to investigate the context |
| | • Panned along the diagonal |
| | • Scaled up and down a cluster of two insets to quickly investigate the visual structure of the patterns |
| | • Continued panning to search for patterns |
| | *Switched to outer-placement view* |
| | • Zoomed out |
| | • Identified new pattern type |
| | • Assessed the original location of the new pattern type through hovering the inset |
| | • Panned to another region |
| | • Investigated the appearance and disappearance of insets based on the zoom level |
| | • Scaled up an inset showing a different pattern type |
| P2 | *Started with inner-placement* |
| | • Obtained an overview through brief zoom out |
| | • Zoomed back in to initial zoom level |
| | • Dragged an aggregate of insets away to study context |
| | • Rapidly panned down the diagonal of the matrix to find dot-like patterns |
| | • Dragged a pile of insets away to study context |
| | • Scaled up and leafed through a pile of insets |
| | • Down scaled a pile of insets as no dot-like patterns were found |
| | • Zoomed out and panned further to gain a better overview |
| | • Found a pile of two insets at the corner of a large square-like, annotated pattern |
| | • Zoomed in to the cluster to inspect the neighborhood of the insets |
| | • Zoomed out again and kept on panning |
| | • Encountered slowness due to a large amount of data loading |
| | • Leafed through a large pile and found some dot-like pattern |
| | • Zoomed into location to explore the local neighborhood |
| | *Switched to outer-placement view* |
| | • Scaled up all the piles of insets to search for dot-like-patterns |
| | • Found one instance and zoomed in to that location |
| | • Rapidly panned to other location while searching for dot-like patterns |
| | *Loaded square-like patterns* |
| | • Explored and found a square-like pattern |
| | • Zoomed in and out investigated the loss of inset |
| | • Zoomed all the way out to look that the square-like patterns globally |
| | • Scaled up and leafed through several piles of insets with unexpected variances |
| P3 | *Started with inner-placement* |
| | • Obtained an overview through gentle panning and zooming |
| | • Adjusted color map of the matrix |
| | • Zoomed out to gain a broader overview |
| | • Zoomed back in to study details |
| | • Dragged a pile of insets to see the nearby context |
| | • Zoomed in to the location containing the insets of the pile |
| | • Zoomed out a bit and panned to other locations |

| | |
|---|---|
| | *Switched to outer-placement view*<br>• Zoomed out a bit and panned to other locations<br>• Investigated disappearance of an inset<br>• Panned to another location<br>• Scaled up a square-like pattern and investigated the detailed visual structure in the inset<br>• Zoomed out while keeping the inset scaled up.<br>• Scaled up another cluster of insets with square-like patterns<br>• Leafed through the cluster to investigate the insets individually as the patterns show some expected details<br>• Panned to another location<br>• Scaled up an inset but kept on panning |
| P4 | *Started with inner-placement*<br>• Panned rapidly down the diagonal<br>• Zoomed out to get a broader overview<br>• Zoomed back in a bit<br>• Dragged a pile of insets away to assess the occluded context<br>• Continued panning to find a dot-like pattern<br>• Scaled up an inset with a dot-like pattern<br>• Zoomed in to the location of the dot-like pattern<br>• Zoomed out to perceive the context at a lower resolution<br>*Switched to outer-placement view*<br>• Moved the mouse cursor over the insets to assess their original location<br>• Zoomed and panned a bit<br>• Tried to combine two insets into a pile<br>• Reset the location of the insets<br>• Scaled up one inset to look at the detailed visual structure<br>• Zoomed into the origin of the scaled-up inset<br>• Investigated when insets disappear<br>• Panned and zoomed further<br>• Zoomed out and compared to distant insets by dragged them next to each other<br>• Zoomed out further out to gain a broader overview |
| P5 | *Started with inner-placement*<br>• Started panning<br>• Identified a specific pattern in an inset and zoomed into the inset's original location<br>• Zoomed out and panned to another location<br>• Found an unexpected pile of insets with "empty" patterns<br>• Zoomed in a bit and confirmed their hypothesis<br>• Zoomed out and found another inset with an expected pattern<br>• Zoomed in to the inset's original location to study the context<br>• Zoomed out and compared the inset to other close-by insets<br>*Switched to outer-placement view*<br>• Scaled up a pile of insets to investigate the detailed visual structure of a square-like pattern<br>• Leafed through the pile of insets<br>• Zoomed into the original location of the pile of insets |

| | |
|---|---|
| | • Scaled up and leafed through other piles of insets in close proximity |
| P6 | *Started with outer-placement*<br>• Gently panned and zoomed to gain an overview<br>• Zoomed in to see more details<br>• Scaled up two insets consecutively<br>• Tried to zoom into the location of an inset via a double click<br>• Zoomed in to the inset's original location using the context menu<br>• Zoomed out again after investigating the context of the annotated pattern<br>• Panned to another location<br>• Scaled up an inset with a square-like pattern<br>*Switched to outer-placement view*<br>• Zoomed in and out to understand the new placement behavior<br>• Zoomed all the way out to find long-distant patterns, i.e., patterns far away from the diagonal of the matrix<br>• Zoomed in to a region without insets<br>• Panned elsewhere along the diagonal<br>• Scaled up an inset showing an unexpected pattern<br>• Zoomed into the inset's original location but stopped half-way to investigate another inset<br>• Panned to another region in the matrix<br>• Closely investigated the detailed visual structure of three insets consecutively |

Table S3: Chronological summary of participant-specific actions and related tasks of the second, qualitative user study with domain experts.

<table>
<tr><th colspan="2">1</th><th>2</th><th>3</th><th>4</th><th>5</th></tr>
<tr><td colspan="6">1. What is your overall impression of the navigation technique? (1: not impressive; 5: very impressive)</td></tr>
<tr><td colspan="2"></td><td></td><td>x</td><td>xxx</td><td>xx</td></tr>
<tr><td colspan="6">2. Do you think the concept of previewing small and aggregated regions as insets within the view is useful for finding interesting regions, i.e., does it shorten the time to decide if we region is worth exploring in detail? (1: disagree; 5: strongly agree)</td></tr>
<tr><td colspan="2"></td><td></td><td></td><td>xx</td><td>xxxx</td></tr>
<tr><td colspan="6">3. How intuitive is the interface? (1: very unintuitive; 5: very intuitive)</td></tr>
<tr><td colspan="2"></td><td>x</td><td></td><td>xxxx</td><td>x</td></tr>
<tr><td colspan="6">4. Are there any parts of the interface that were not intuitive?</td></tr>
<tr><td colspan="6">• With a very brief introduction (<5min) things were understandable<br>• Re-sync button not obvious<br>• Border gallery<br>• Fading lines are confusing<br>• Pile aggregation type not obvious<br>• Disappearance of snippets is confusing</td></tr>
<tr><td colspan="6">5. How useful is the tool in its current form to you? (1: not useful at all; 5: very useful)</td></tr>
<tr><td colspan="2"></td><td>x</td><td>xx</td><td>xx</td><td>x</td></tr>
<tr><td colspan="6">6. Is this kind of exploration currently possible in any other form? (1: not at all; 5: very similar tools exist)</td></tr>
<tr><td colspan="2">xxx</td><td>x</td><td>xx</td><td></td><td></td></tr>
<tr><td colspan="6">7. If so, how does our tool compare against the other methods in terms of performance (A) and features (B)? (1: worse; 5: much better)</td></tr>
<tr><td>A</td><td>x</td><td>x</td><td></td><td>x</td><td></td></tr>
<tr><td>B</td><td></td><td>x</td><td>x</td><td>x</td><td></td></tr>
<tr><td colspan="6">8. Which (navigation) features (if any) are missing that would make this tool more useful? (Please sort by importance)</td></tr>
<tr><td colspan="6">• User-resizable snippets (2x)<br>• Change color scale<br>• Scale bar for snippet size<br>• Delete snippets<br>• Add new snippets manually (2x)<br>• Toggle between placing techniques<br>• Pin snippets<br>• Adjust aggregate representation<br>• Filter displayed snippets by some value</td></tr>
<tr><td colspan="6">9. Imagine all missing features are implemented, how useful would do you think would this tool be to the research community? (1: not useful at all; 5: very useful)</td></tr>
<tr><td colspan="2"></td><td></td><td></td><td>xx</td><td>xxxx</td></tr>
<tr><td colspan="6">10. Could you imagine this tool to be useful for other applications like matrices / heatmaps, geographic maps, large images / high content screens? (1: not at all; 5: very much)</td></tr>
<tr><td colspan="2"></td><td></td><td></td><td>x</td><td>xxxxx</td></tr>
</table>

Table S4: Closing questionnaire of the second user study. Each "x" stands for the answer of one of the six participants. The lists for question 4 and 8 have been paraphrased and combined for brevity. We employed end labeling, i.e., only points 1 and 5 were labeled as shown above. All questions were optional.

| Parameter | INSIDE | OUTSIDE |
|---|---|---|
| insets.labelPosition | hidden | |
| insets.minSize | 32 | |
| insets.maxSize | 56 | |
| insets.sizeStepSize | 2 | |
| insets.scale | 1 | |
| insets.scaleSizeBy | size | |
| insets.additionalZoom | 1 | |
| insets.onClickScale | 3 | |
| insets.fill | white | |
| insets.fillOpacity | 1 | |
| insets.borderColor | white | |
| insets.borderWidth | 1 | |
| insets.borderOpacity | 1 | |
| insets.borderRadius | 4 | |
| insets.leaderLineColor | white | |
| insets.leaderLineWidth | 2 | |
| insets.leaderLineOpacity | 1 | |
| insets.leaderLineStubWidthMin | | 2 |
| insets.leaderLineStubWidthMax | | 4 |
| insets.leaderLineStubLength | | 12 |
| insets.leaderLineDynamic | | true |
| insets.leaderLineDynamicMinDist | | 24 |
| insets.leaderLineDynamicMaxDist | | 120 |
| insets.dropDistance | 1 | |
| insets.dropBlur | 3 | |
| insets.dropOpacity | 0.8 | |
| insets.opacity | 1 | |
| insets.showClusterSize | true | |
| insets.isDraggingEnabled | true | false |
| insets.loadHiResOnScaleUp | true | |
| insets.isImgSelectable | false for PATTERN and true for TYPE | |
| meta.insetThreshold | 12 | |
| meta.gridSize | 72 | 52 |
| meta.minClusterSize | 3 | |
| meta.maxClusterDiameter | 96 | 64 |
| meta.boostContext | 1 | 0 |
| meta.boostDetails | 1 | 10 |
| meta.boostLocality | 1 | 1 |
| meta.boostLayout | 1 | 5 |
| meta.cooling | 1 | |
| meta.reheat | 0.05 | |

Table S5: Parameter settings for INSIDE and OUTSIDE of the controlled user study with gigapixel images. The settings prefixed with "insets" refer to `Insets2dTrack` and the settings prefixed with "meta" refer to `AnnotationsToInsetsMetaTrack`. See `https://github.com/flekschas/higlass-scalable-insets` for more information.

Table S6: Aggregation of the absolute votes from the closing questionnaires (Table S2 and S4). Mean values are shaded by their values. *Note that Q3 of study 1 was answered by 17 out of 18 and Q7 of study 2 was answered by 3 out of 6 participants only. Regarding the latter two, the *missing* votes are due to the fact that 3 participants for Q6 voted that no other tool currently supports exploration like Scalable Insets and, hence, did not have to answer Q7.

| | **Study 1** | | | **Study 2** | | | | | | | | |
|---|---|---|---|---|---|---|---|---|---|---|---|---|
| | **Q1** | **Q2** | **Q3*** | **Q1** | **Q2** | **Q3** | **Q5** | **Q6** | **Q7A*** | **Q7B*** | **Q9** | **Q10** |
| **Mean** | **3.9** | **3.8** | **4.5** | **4.2** | **4.7** | **3.7** | 3.5 | 1.8 | 2.3 | 3.0 | **4.7** | **4.8** |
| **1, 2, and 3** | 5 | 5 | 3 | 1 | 0 | 1 | 3 | 6 | 2 | 2 | 0 | 0 |
| **4 and 5** | 13 | 13 | 14 | 5 | 6 | 5 | 3 | 0 | 1 | 1 | 6 | 6 |

Table S7: Technical setup of both user studies

| | |
|---|---|
| **Computer** | Apple MacBook Pro 2016 |
| **Processor** | Intel 2.7 GHz quad-core |
| **Memory** | 16 GB |
| **Display** | 15" |
| **Effective resolution** | 1440 × 900 pixels |
| **Operating system** | macOS 10.12 |
| **Input device** | Standard two-button mouse with a scroll wheel |

### Supplementary Pseudo Code

```
input  : insets I, maximum inset size size, view height height, and view width width
numBinsX ← ⌊width/size⌋;
numBinsY ← ⌊height/size⌋;
binsLeft = binsRight = list of length numBinsY;
binsTop = binsBottom = list of length numBinsX;
for inset i ∈ I do
    BinX ← ⌊i.x/(width/numBinsX)⌋;
    BinY ← ⌊i.y/(height/numBinsY)⌋;
    dLeft ← i.x + GetBinValue(binsLeft, BinY) ∗ size;
    dRight ← width − i.x + GetBinValue(binsRight, BinY)⌋) ∗ size;
    dTop ← i.y + GetBinValue(binsTop, BinX) ∗ size;
    dBottom ← height − i.y + GetBinValue(binsBottom, BinX) ∗ size;
    closestSide ← i.side or GetClosestSide (dLeft, dRight, dTop, dBottom);
    switch closestSide do
        case left do
            i.side ← left;
            IncrementBinValue (binsLeft, BinY));
        end
        case right do
            i.side ← right;
            IncrementBinValue (binsRight, BinY));
        end
        case top do
            i.side ← top;
            IncrementBinValue (binsTop, BinX));
        end
        case bottom do
            i.side ← bottom;
            IncrementBinValue (binsBottom, BinX));
        end
    end
end
```

**Algorithm S1: Gallery Layout: Assign Insets to a Side**. Insets are initially assigned to the closest side with the fewest number of insets already assigned to the bin of the corresponding x or y screen location of the inset. Upon pan or zoom, the side is not changed anymore to provide a stable map of the insets' positions.

```
input  : insets I, maximum area maxA, and maximum distance maxD
output : clusters C

C ← empty list;
for inset i ∈ I do
    d ← maxD;
    closest ← NIL;
    for cluster c ∈ C do
        if AreaOf (c) > maxA then
            continue
        end
        distance ← L2Dist(i, c);
        if distance < d then
            closest ← c;
            d ← distance;
        end
    end
    if IsNotNil(closest) and CombinedAreaOf(i, c) ≤ maxA then
        AddToCluster(c, i);
    end
    else AppendToList (C, CreateNewCluster(i)) ;
end
return C
```

**Algorithm S2: Initial clustering**. `AreaOf` computes the area of the convex hull defined by a cluster of points. Additionally, `CombinedAreaOf` computes the area of the convex hull defined by a cluster that is extended by a point. `L2Dist` computes the Euclidean distance between an inset and the center of a cluster in pixel coordinates. Details about `AddToCluster` are provided in Algorithm S3.

```
input   :cluster c, inset iNew
d ← Infinity;
closest ← NIL;
for inset i ∈ C.I do
    if C.FNN = NIL then
        C.FNN ← empty priority queue in descending order;
    end
    distance ← L2Dist (i, iNew);
    if distance < d then
        closest ← i;
        d ← distance;
    end
end
AddToSet (C.I, i);
AddToQueue (C.FNN, (d, closest, iNew));
```

**Algorithm S3: Add an Inset to a Cluster**. In order to quickly determine breakpoints of a cluster, we are keeping track of the nearest neighbor of insets as they are being added to a cluster. Upon zooming in we then only have to evaluate if the distance of the farthest nearest neighbor is larger than the threshold and if it is we split the cluster into two separate clusters such that the first cluster contains all patterns closer to the first pattern of the pair of farthest nearest neighbors and the second cluster is composed of all the other patterns. E.g., Figure 5.5.i shows the farthest nearest neighbor pair with a black dashed line.

```
input  : clusters C, zoom direction z, maximum distance maxD, maximum area maxA
output : updated clusters C

if z == 1 then
    maxD = maxD × 1.5;
    for cluster c ∈ C do
        if SizeOf (c) > 1 then
            fnn ← GetFNN (c);
            distance ← L2Norm (fnn);
            if distance > maxD then
                [c1,c2 ] ← SplitClusterAt (c, fnn);
                AppendTo (C, [c1,c2 ]);
            end
        end
    end
else if z == −1 then
    maxD = maxD × 0.5;
    for cluster c1 ∈ C do
        for cluster c2 ∈ C do
            if c1 ≠ c2 and L2Dist(c1, c2) ≤ maxD and CombinedAreaOf(c1, c2) ≤
              maxA then
                MergeClusters (c1, c2);
            end
        end
    end
```

**Algorithm S4: Update clustering**. Upon zooming in or out, clusters are evaluated whether they need to be split or merged. On zooming in (z == 1), split a cluster if the L2 distance of the farthest nearest neighbors (`FNN`) is larger than 1.5 × `maxD`. The `FNN` are defined when a point is added to a cluster. The distance between the nearest neighbor and the newly added point is stored. `FNN` is then defined as the pair of points on the cluster that was farthest away. Internally, `FNN` is a cached property of each cluster instance to quickly determine if and where a cluster would need to be split. On zooming out (z == -1), the distance between all pairs of clusters are compared. If the distance between two clusters `c1` and `c2` is smaller than 0.5 × `maxD` and the combined are is smaller than `maxA` the clusters are merged.

**input** : cluster c, importance property p
**output** : representative insets R

```
if p ≠ NIL then
    r1 ← GetMostImportant (C.I,p);
end
else r1 ← GetLargest (C.I);
;
r2 ← GetClosestToCenterExcept (C.I, C.centroid, [r1]);
r3 ← GetFarthestFromCenterExcept (C.I, C.centroid, [r1,r2]);
r4 ← GetFarthestFromInsetExcept (C.I, r3, [r1,r2,r3]);
return [r1,r2,r3,r4]
```

**Algorithm S5: Gallery Representatives**. Heuristic for choosing the representative insets of an aggregated gallery inset. `GetClosestToCenterExcept`, `GetFarthestFromCenterExcept`, and `GetFarthestFromInsetExcept` operate in Euclidean space.
